# Combining Cell-Intrinsic and -Extrinsic Resistance to HIV-1 By Engineering Hematopoietic Stem Cells for CCR5 Knockout and B Cell Secretion of Therapeutic Antibodies

**DOI:** 10.1101/2024.03.08.583956

**Authors:** William N. Feist, Sofia E. Luna, Kaya Ben-Efraim, Maria V. Filsinger Interrante, Nelson A. Amorin, Nicole M. Johnston, Theodora U. J. Bruun, Hana Y. Ghanim, Benjamin J. Lesch, Amanda M. Dudek, Matthew H. Porteus

## Abstract

Autologous transplantation of *CCR5* null hematopoietic stem and progenitor cells (HSPCs) is the only known cure for HIV-1 infection. However, this treatment is limited because of the rarity of *CCR5*-null matched donors, the morbidities associated with allogeneic transplantation, and the prevalence of HIV-1 strains resistant to CCR5 knockout (KO) alone. Here, we propose a one-time therapy through autologous transplantation of HSPCs genetically engineered *ex vivo* to produce both CCR5 KO cells and long-term secretion of potent HIV-1 inhibiting antibodies from B cell progeny. CRISPR-Cas9-engineered HSPCs maintain engraftment capacity and multi-lineage potential *in vivo* and can be engineered to express multiple antibodies simultaneously. Human B cells engineered to express each antibody secrete neutralizing concentrations capable of inhibiting HIV-1 pseudovirus infection *in vitro*. This work lays the groundwork for a potential one-time functional cure for HIV-1 through combining the long-term delivery of therapeutic antibodies against HIV-1 and the known efficacy of *CCR5* KO HSPC transplantation.

## Introduction

There are 39 million people worldwide living with HIV infection.^1^ While great advances in small-molecule antiretroviral therapy (ART) have resulted in combination drug regimens that can control viral loads, in 2022 alone there were over 600,000 AIDS-related deaths, underlining that HIV remains a global epidemic with substantial morbidity and mortality.^1–3^ This is due in part to the strict adherence necessary for ART to remain effective.^4–6^ Current regimens call for lifetime daily dosing to suppress long-term viral reservoirs and prevent viral rebound.^7,8^ Because of this, significant research efforts have sought to identify novel treatment modalities that provide a more durable long-term cure.

The discovery of highly potent HIV-1 inhibiting antibodies with relatively long half-lives has led to numerous clinical trials for sustained control of HIV-1.^9,10^ The first monoclonal antibody clinically approved to treat HIV-1, Ibalizumab, acts as a post-attachment inhibitor by binding HIV-1’s primary receptor on human cells, CD4.^11–14^ In addition to this, numerous extremely potent antibodies that act against diverse HIV-1 subtypes have been identified to bind and neutralize HIV-1 directly.^15,16^ These antibodies, known as broadly neutralizing antibodies (bNAbs), target highly conserved regions of the viral envelope to inhibit infection. Recent and ongoing trials are testing the efficacy of targeting multiple env epitopes simultaneously to prevent the generation of resistance mutations.^9,10,17–19^ These trials have shown that bNAbs can maintain viral suppression and prevent the formation of escape mutants so long as antibody titers remain above a therapeutic threshold. While the longer half-life of antibodies does allow for reduced frequency of dosing, lifelong repeated administration would still be required to maintain efficacy.

One promising method for sustained delivery of bNAbs is through AAV-mediated delivery of antibody expression cassettes, known as vectored immunoprophylaxis.^20–25^ Trials in mice, macaques, and recently humans have demonstrated the potential of this strategy for sustained secretion of therapeutic anti-HIV-1 antibodies. However, pre-existing immunity against AAV vectors, low levels of antibody secretion, uncertainty regarding long-term expression, and seroconversion preventing re-dosing remain challenges limiting this approach.^26,27^ Another potential method for long-term maintenance of antibody expression is the direct editing of autologous B cells. B cells can be directly engineered for custom antibody expression from the B cell receptor locus, allowing for participation of the custom antibody in the humoral immune response.^28–32^ However, there is currently no clinical protocol for the engraftment of B cells in patients and it remains unclear how long they may persist in the body.

Hematopoietic stem cell transplantation (HSCT) of *CCR5* knockout (KO) cells is the only reported long-term functional cure for HIV-1 infection. Demonstrated first in the cases of the Berlin and London patients, allogeneic HSCT with cells from donors carrying the naturally occurring *CCR5*-*Δ32* KO mutation resulted in reconstitution with cells that were resistant to their CCR5-tropic (R5-tropic) HIV-1.^33,34^ After treatment interruption of ART, each patient maintained viral suppression and was considered functionally cured of their infection. While this strategy has shown continued success, it is not widely available to the vast majority of patients.^35^ The rarity of identifying a *CCR5* KO matched donor and the morbidities associated with allogeneic transplantation, such as graft versus host disease (GvHD), limit this treatment to only a small fraction of patients who require HSCT for an underlying malignancy.^36–38^ Moreover, HSCT with *CCR5* KO cells is ineffective in patients carrying CXCR4-tropic (X4-tropic) HIV-1 strains that do not rely on CCR5 for cellular entry. This was demonstrated in the case of the Essen patient, where *CCR5* KO HSCT followed by ART interruption resulted in rebound of X4-tropic virus.^39^ X4-tropic HIV-1 is estimated to be present in 18% to 52% of patients, highlighting the need for strategies to combat these strains.^40,41^

Autologous HSCT with genetically modified cells is a promising strategy to overcome the risk of GvHD and the need for rare donor cells. One recent trial demonstrated the feasibility of autologous transplantation of cells genetically modified for *CCR5* KO in a patient with HIV-1.^42^ However, low editing rates resulted in a failure to prevent viral rebound, underscoring the importance of efficient modification to minimize viral replication in unedited cells. Other strategies have modified hematopoietic stem and progenitor cells (HSPCs) with *CCR5* KO, lentiviral delivery of HIV-1 inhibiting proteins, neutralizing antibodies, and inhibitory RNAs against *CCR5* and viral targets.^42–48^ Several of these strategies benefit from layering multiple methods of HIV-1 inhibition to control both R5-tropic and X4-tropic HIV-1, however, limited editing efficiency and the risk of malignancy resulting from insertional mutagenesis have hampered their application.^49^ Meanwhile, advances in precision editing with CRISPR-Cas9 have allowed for high-efficiency cellular engineering without the risks associated with lentiviral integration. Recently, we reported a CRISPR-Cas9-based strategy in HSPCs for simultaneous KO out of *CCR5* with knock-in of expression cassettes for two HIV-1 inhibiting proteins.^50^ While this work demonstrated the feasibility of high-efficiency editing to deliver multilayered genetic resistance to HIV-1, it relies on a cell-autonomous resistance scheme that leaves unedited cells vulnerable to infection. As such, we seek to develop a strategy that can deliver both cell-intrinsic and -extrinsic protection to inhibit infection in both edited and unedited cells.

Here, we seek to combine the success of *CCR5* KO HSCT with the potency of sustained antibody therapy through a simultaneous knockout knock-in gene editing strategy in HSPCs. Our system is designed to deliver broad resistance against CCR5-tropic HIV-1 through *CCR5* KO and to include additional layers of non-cell-autonomous protection through the secretion of multiple HIV-1 inhibiting antibodies that act broadly against both R5- and X4-tropic viruses. To combat the formation of escape mutants, we designed our system for use with multiple well-established antibodies targeting diverse epitopes. In this study, we demonstrate that HSPCs can be efficiently edited at the *CCR5* locus for knock-in of multiple HIV-1 inhibiting antibody expression cassettes. Importantly, these engineered HSPCs maintain their engraftment capacity and multilineage potential following transplantation in immunodeficient mice. Moreover, we show that primary human B cells carrying the antibody expression cassettes secrete neutralizing levels of antibodies *in vitro*. Overall, this work establishes a strategy for autologous transplantation of HSPCs modified for long-term secretion of therapeutic antibodies from B cell progeny.

## Results

### HIV-1 inhibiting antibodies maintain function with a peptide linker

Recent trials investigating direct injection of anti-HIV-1 antibodies to control disease highlight the need to use multiple antibodies targeting different epitopes in combination to prevent viral escape.^17–19^ To this end, we utilize well-established antibodies including the clinically approved antibody Ibalizumab and several bNAbs that target different highly conserved regions of the HIV-1 envelope: 10-1074 (V3 Loop), PGDM1400 (V1/2 Loop), CAP256V2LS (V1/2 Loop), 3BNC117 (CD4 binding site), and 1-18 (CD4 binding site).^51–55^

While traditional production of these IgG antibodies involves transfection of cell culture with two separate plasmids encoding the IgG heavy and light chain, our delivery system necessitates the antibody components be delivered as a single transcript for knock-in to the *CCR5* locus. However, expression from B cell progeny that are producing antibodies from the native immunoglobulin loci may result in mispairing of therapeutic antibodies with endogenous heavy or light chains, resulting in dysfunctional epitope recognition. For this reason, we employed a linker system to physically pair the therapeutic antibody heavy and light chain to mitigate the risk for mispairing and the potential formation of deleterious antibody products (Fig. 1a).^56^

**Figure 1:**
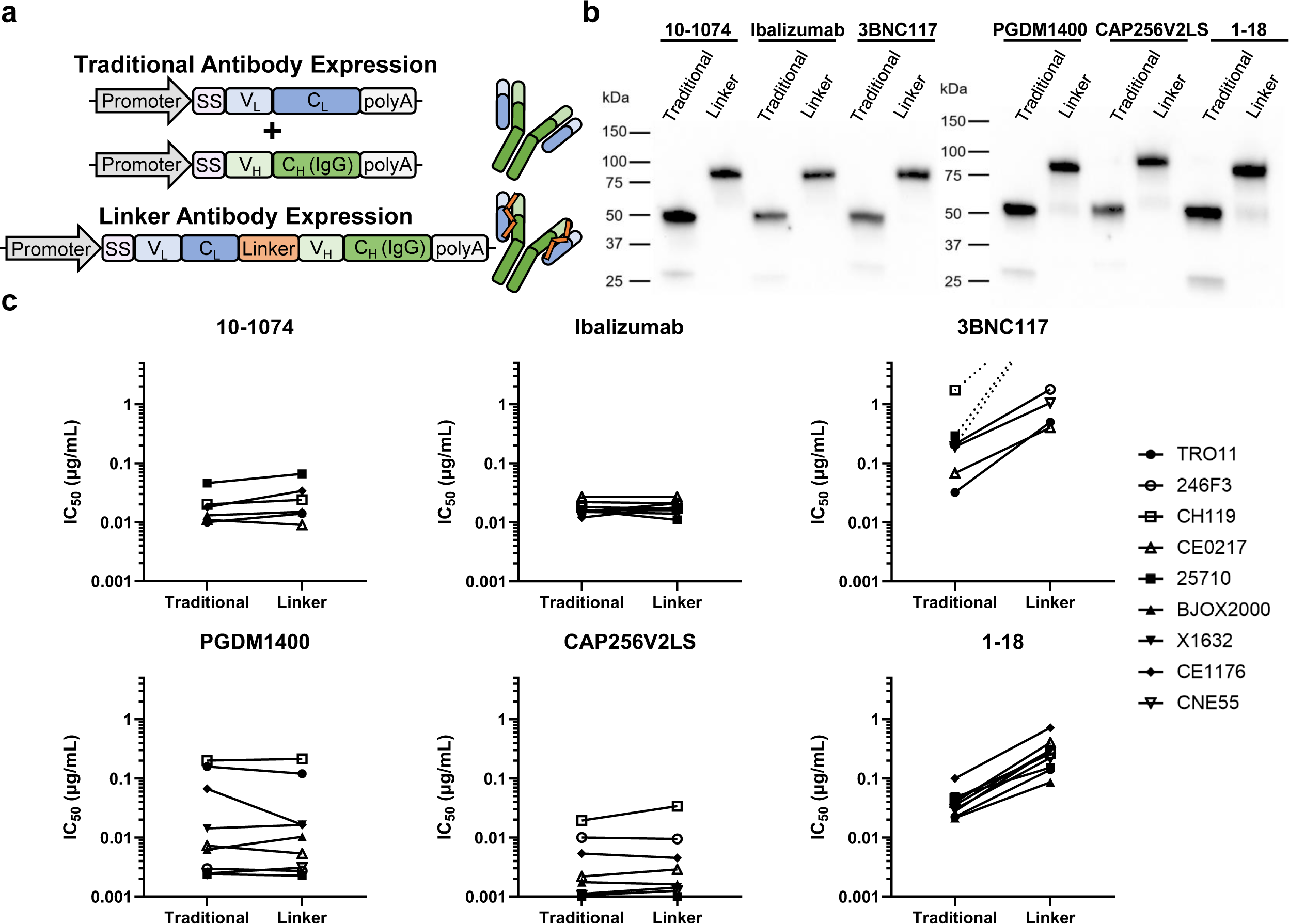
HIV inhibiting antibodies maintain function with a peptide linker. **a)** Diagram of antibody expression with and without a peptide linker. **b)** SDS-PAGE western blot of purified traditional and linker antibodies. **c)** IC_50_ of traditional and linker antibodies against a panel of HIV-1 pseudoviruses measured *in vitro* with TZM-bl infection assay (IC_50_ was calculated from technical duplicate infections across serial dilutions of each antibody). Pseudoviruses not shown on a graph were not inhibited within the tested antibody concentrations (≤ 5 µg/mL).

To determine if the addition of the linker impacts function, each antibody was produced in the conventional manner from a separate cassette for the light and heavy chains (traditional antibody) and from a single cassette with a linker pairing the light and heavy chains (linker antibody) (Fig. 1a). Purified antibodies were applied to western blot analysis under reducing conditions and stained for IgG. Traditional antibodies were detected at the expected mass for a heavy chain alone at approximately 50 kDa, while linker antibodies formed a band with an expected higher molecular weight at 80 kDa, corresponding to the increased mass of the linker and light chain paired with the heavy chain (Fig. 1b). Each antibody was then tested against a panel of HIV-1 pseudoviruses representing the global diversity of HIV-1 in a TZM-bl infection assay to ensure neutralization potency was preserved with the added linker.^57^ The traditional IgG and linker versions of 10-1074, Ibalizumab, PGDM1400, and CAP256V2LS show comparable efficacy as demonstrated by measured half-maximal inhibitory concentration (IC_50_) for each pseudovirus (Fig. 1c and Supplementary Fig. 1). Two antibodies, 3BNC117 and 1-18, demonstrated a marked reduction in neutralization efficacy when expressed with a peptide linker. Interestingly, these two antibodies both target the CD4 binding site on HIV-1, indicating that further optimization may be needed to employ a linker for this subset of antibodies. Overall, these results demonstrate that HIV-1 inhibiting antibodies can maintain function when expressed from a single transcript with a peptide linker physically pairing the light and heavy chains.

### Simultaneous *CCR5* KO and Antibody Cassette Knock-in in CD34^+^ HSPCs

The modification of CD34^+^ HSPCs is a promising strategy to deliver long-term HIV-1 treatment through hematopoietic reconstitution with resistant cells. However, high-efficiency editing is needed to mitigate the potential for escape mutants to form in non-edited cells and to deliver therapeutically relevant antibody levels. To this end, we employ CRISPR-Cas9 editing with electroporation of ribonucleoprotein (RNP) and adeno-associated virus serotype 6 (AAV6) delivery of DNA donor templates for knock-in by homology directed repair (HDR). *CCR5* KO is achieved with a highly efficient single guide RNA (sgRNA) we previously demonstrated to induce KO INDELs (Supplementary Fig. 2a) that confer resistance to CCR5-tropic HIV-1 infection in primary CD4^+^ T cells.^50^ Off-target analysis previously performed for this sgRNA showed it to be specific with no measurable off-target INDEL formation.^50^ AAV6-delivered donor templates for each linker antibody contain homology flanking the *CCR5* cut site and are driven by a previously defined B cell promoter, the EEK promoter (Fig. 2a), for strong expression in B cells as the professional antibody secreting cell type.^45^ Cord blood (CB) CD34^+^ HSPCs were edited with each antibody construct individually. We also edited cells with 10-1074 and Ibalizumab constructs in combination as proof of concept for simultaneous delivery of multiple antibodies. Knock-in analysis through in-out droplet digital PCR (ddPCR) determined that each antibody construct individually was incorporated in an average of 31% to 41% of alleles (Fig. 2b and Supplementary Fig. 2b). When used in combination, 10-1074 and Ibalizumab knock-in was detected in up to 44% of alleles. We designed a construct-specific ddPCR assay to determine the prevalence of each antibody within the bulk cell population and found slightly higher knock-in of the Ibalizumab cassette compared to 10-1074 (Fig. 2c and Supplementary Fig. 2b-e). While we focus on the use of 10-1074 and Ibalizumab together, further combinations can be made through editing with any two or three antibody cassettes simultaneously in CD34^+^ HSPCs (Supplementary Fig. S3).

**Figure 2:**
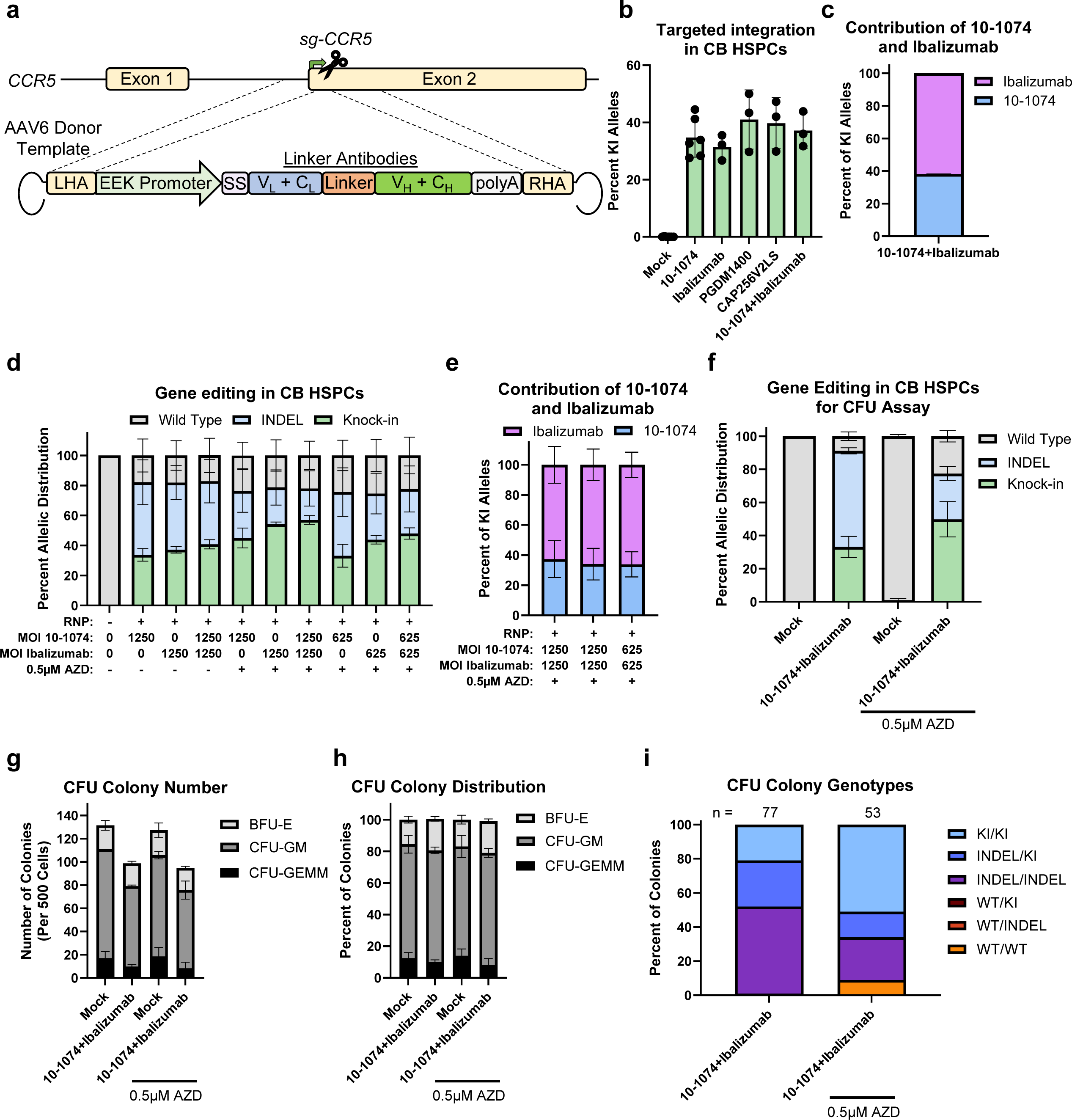
Efficient targeted integration of antibody expression cassettes at the *CCR5* locus in HSPCs. **a)** Schematic of gene editing strategy to deliver linker antibody expression cassettes to the *CCR5* locus. **b)** Allelic integration frequency in CB CD34^+^ HSPCs targeted with AAV6 cassettes for linker antibodies as indicated. Each AAV6 was used at a multiplicity of infection (MOI) of 1250. Values represent independent biologic donors with n = 6 for mock and 10-1074 conditions and n = 3 for Ibalizumab, PGDM1400, CAP256V2LS, and 10-1074+Ibalizumab conditions. **c)** Percent of KI alleles integrated with 10-1074 or Ibalizumab within the “10-1074+Ibalizumab” targeted cells shown in b (n = 3). **d)** Distribution frequency of WT, INDEL, and KI alleles in CB CD34^+^ HSPCs targeted at *CCR5* with the AAV6 MOI as indicated with or without 0.5 μM AZD7648 (n = 3). **e)** Percent of knock-in alleles integrated with 10-1074 or Ibalizumab within the “10-1074+Ibalizumab” targeted cells shown in d (n = 3). **f)** Distribution frequency of WT, INDEL, and KI alleles in CB CD34^+^ HSPCs used for the CFU assay described in G, H, and I. Cell were targeted at *CCR5* with an AAV6 MOI of 625 for each construct with or without 0.5 µM AZD7648 (n = 2). **g)** Number of BFU-E, CFU-GM, and CFU-GEMM colonies formed per 500 cells plated in the CFU assay (n = 2, with technical duplicates wells for each donor). **h)** Relative frequency of BFU-E, CFU-GM, and CFU-GEMM colonies formed within the CFU assay (n = 2, with technical duplicate wells for each donor). **i)** Frequency of genotypes from single-cell colonies within the “10-1074+Ibalizumab” targeted condition treated with or without 0.5 µM AZD7648 (n represents the total number of colonies genotyped across both biological donors). All replicates represent independent biological donors unless otherwise noted. All bars represent mean and error bars represent standard deviation (SD).

### Small molecule inhibition of NHEJ improves knock-in efficiency of antibody constructs

Inhibition of NHEJ by inhibiting DNA-PKcs with the small molecule AZD7648 has recently been shown to improve allelic knock-in frequency in RNP/AAV6 HDR-based editing.^58^ Therefore, we tested the use of AZD7648 treatment with our constructs to increase editing frequency and reduce the multiplicity of infection (MOI) of AAV6 used for targeted knock-in. We found that AZD7648 treatment allowed us to reduce the AAV6 used by half while maintaining a high frequency of knock-in and *CCR5* KO resulting from either knock-in or INDEL formation (Fig. 2d). NHEJ inhibitor treatment did not impact the relative prevalence of 10-1074 and Ibalizumab in cells edited with both constructs (Fig. 2e).

To analyze the impact of AZD7648 treatment on differentiation capacity and cell health, we performed colony forming unit (CFU) assay analysis that measures myelo-erythroid differentiation potential *in vitro*. Two donors of CB CD34^+^ cells were treated with or without AZD7648 and received electroporation only (mock) or gene editing using a low MOI of both 10-1074 and Ibalizumab AAV6 together. As expected, we found that AZD7648 treatment increased knock-in frequency in the bulk cell population (Fig. 2f). AZD7648 treatment did not impact total colony formation or the distribution of colony formation between the three major sub-types, CFU-GEMM (colony forming unit-granulocyte, erythroid, macrophage, megakaryocyte), CFU-GM (colony forming unit-granulocyte and monocyte), and BFU-E (burst forming unit-erythroid) (Fig. 2g-h). Single colonies were genotyped to determine the impact of AZD7648 on mono-allelic and bi-allelic knock-in frequency, as well as INDEL formation. We found that AZD7648 treatment resulted in a 2.4-fold increase in the proportion of bi-allelic knock-in events across colony sub-types (21% to 51%) (Fig. 2i). Indeed, just 9% of colonies maintained a wild-type (WT) allele following treatment with AZD7648. These results demonstrate that we can achieve efficient *CCR5* KO and knock-in of our antibody expression cassettes in CD34^+^ HSPCs, with further improved knock-in frequencies through the use of small molecule inhibition of NHEJ.

### Engineered HSPCs maintain engraftment capacity and multilineage reconstitution in immunodeficient mice

Genetically engineered HSPCs must maintain the ability to engraft and reconstitute the hematopoietic lineages in order to remain a viable therapeutic modality for long-term HIV-1 treatment. We therefore edited CB CD34^+^ cells with 10-1074, Ibalizumab, or both 10-1074 and Ibalizumab together (at higher AAV6 MOI and without AZD7648) and transplanted them in sub-lethally irradiated immunodeficient NSG mice (Fig. 3a). Prior to transplantation, we determined that each edited condition had similar frequencies of knock-in (33-38%) and *CCR5* KO INDEL formation (52-56%), with a slight prevalence of Ibalizumab knock-in in the combination edited cells (Fig. 3b-c). Engraftment, multilineage reconstitution, and editing frequencies of transplanted cells were measured at 16 weeks post-transplantation. Human chimerism was measured in bone marrow and spleen through flow cytometry analysis of human and mouse markers. Cells from each condition successfully engrafted in the bone marrow and showed migration to the spleen, a secondary lymphatic site, with similar human chimerism between each gene-edited condition (Fig. 3d-e). The observed decrease in total human engraftment due to RNP/AAV6-based editing was in line with previously published works.^50,59–62^ Hematopoietic lineage distribution was consistent between conditions, with each editing condition producing similar frequencies of CD19^+^ B lineage cells and CD33^+^ myeloid lineage cells in both the bone marrow and spleen, indicating that our editing strategy does not introduce reconstitution bias (Fig. 3f-g). Knock-in alleles were present at the endpoint in both the bone marrow and spleen, though with a lower knock-in frequency than the bulk cells at the time of engraftment (Fig. 3h). Importantly, isolated CD19^+^ cells maintained knock-in alleles, indicating that the introduction of exogenous antibody cassettes does not inhibit B cell formation *in vivo* (Fig. 3i). *CCR5* KO alleles were also maintained in all conditions receiving RNP (Fig. 3j and Supplementary Fig. 4a). These analyses demonstrate that our engineered HSPCs maintain their ability to engraft and reconstitute the hematopoietic lineages, providing promise for the development of an autologous transplant therapy. However, the relatively low frequency of integrated alleles in the cells at the engraftment endpoint warranted further optimization. The results shown in Fig. 2 demonstrated that an approach to optimization was to incorporate DNA-PKcs inhibition while lowering the MOI of AAV6.

**Figure 3:**
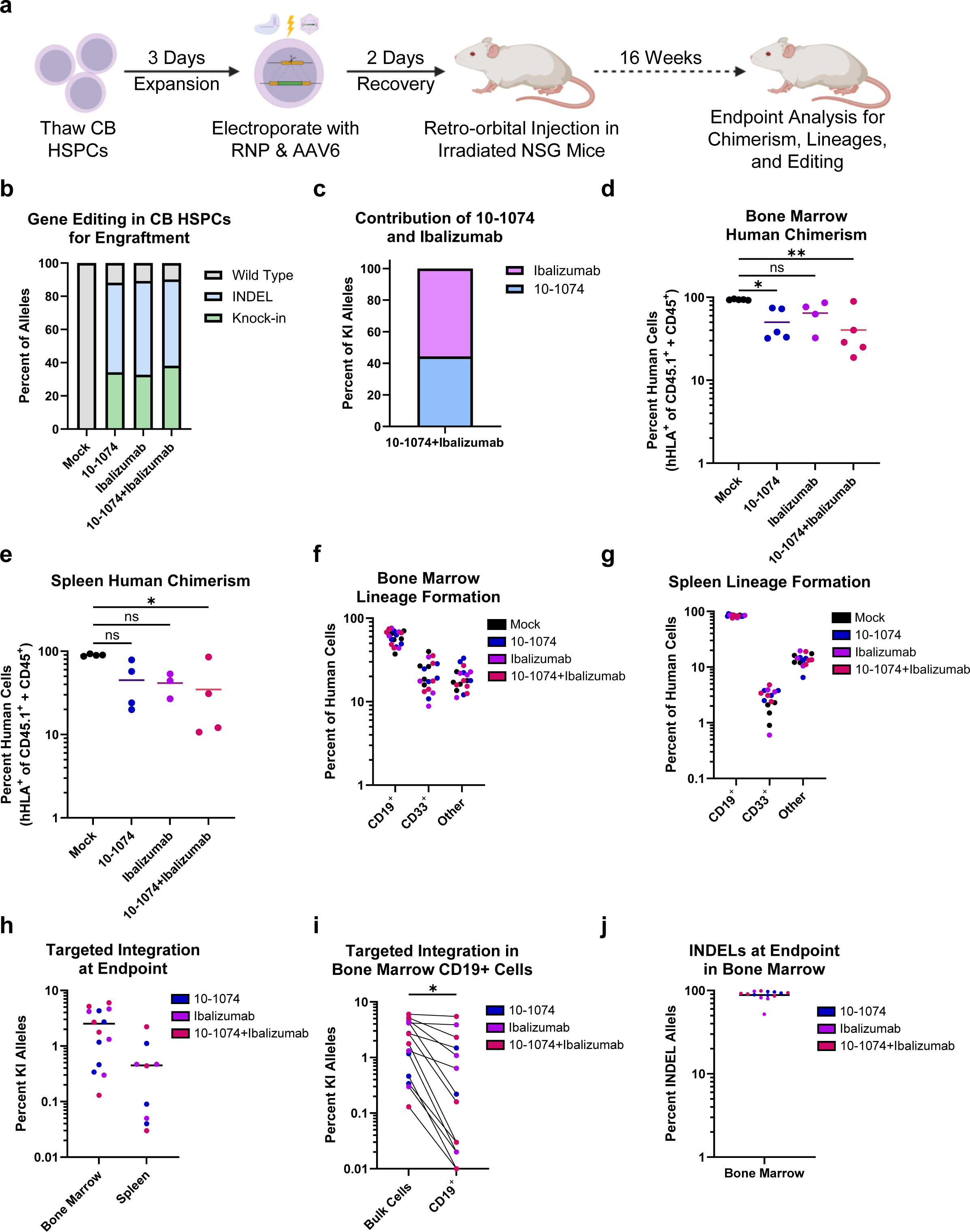
Antibody edited HSPCs maintain engraftment capacity and multilineage reconstitution *in vivo*. **a)** Diagram for editing and engraftment of CB CD34^+^ HSPCs in NSG mice to characterize engraftment capacity and multilineage potential. **b)** Distribution frequency of WT, INDEL, and KI alleles in CB CD34^+^ HSPCs prior to transplantation in NSG mice. Cells were targeted at *CCR5* with an AAV6 MOI of 1250 for each antibody construct (n = 1 pooled sample from 5 donors). **c)** Percent of KI alleles integrated with 10-1074 or Ibalizumab within the 10-1074+Ibalizumab targeted cells shown in panel b (n = 1). **d)** Percent human cell chimerism in the bone marrow at 16 weeks post-transplantation (n = 5 for mock, 10-1074, and 10-1074+Ibalizumab, n = 4 for Ibalizumab). One-way ANOVA Kruskal-Wallis test plus Dunn’s multiple comparisons test (ns, not significant, P = 0.3287; *P = 0.0496; **P = 0.0082). **e)** Percent human cell chimerism in the spleen at 16 weeks post-transplantation (n = 4 for mock, 10-1074, and 10-1074+Ibalizumab, n = 3 for Ibalizumab). One-way ANOVA Kruskal-Wallis test plus Dunn’s multiple comparisons test (ns, not significant, P = 0.0806 for Mock vs 10-1074, P = 0.1363 for Mock vs Ibalizumab; *P = 0.0216). **f)** Percent of human cells in the bone marrow (n are the same as in d) or **g)** spleen (n are the same as in e) that are CD19^+^ (B cell lineage), CD33^+^ (myeloid cell lineage), or within other lineages in mice engrafted with mock (black) or gene edited HSPCs (10-1074, blue; Ibalizumab, purple; 10-1074 and Ibalizumab, red). **h)** Percent of human alleles from the bone marrow or spleen with knock-in of the indicated antibody constructs (n are the same as in d and e for bone marrow and spleen, respectively). **i)** Percent of human alleles with knock-in from the bulk bone marrow (as shown in panel h) or in positively selected bone marrow CD19^+^ cells (n are the same as in d). Lines connect dots representing measurements from the same mice. Two-tailed Mann-Whitney test (*P = 0.0135). **j)** Percent of human alleles from the bone marrow with an INDEL at *CCR5* (n are the same as in d). This analysis does not include alleles with KI. All bars represent mean. All dots represent individual mice engrafted with mock HSPCs or HSPCs edited with AAV6 and RNP for the antibody construct(s) indicated.

### High-efficiency editing in CD34^+^ HSPCs allows for maintenance of integrated alleles following engraftment *in vivo*

Achieving a high frequency of integrated antibody cassettes in engrafted cells will be important for future studies determining the minimum editing threshold needed to achieve clinically relevant concentrations of each antibody. To this end, we employed AZD7648 treatment to improve the initial knock-in frequency in the CD34^+^ HSPCs prior to transplantation. Additionally, the use of AZD7684 allowed us to employ reduced AAV6 doses to potentially improve cell health and engraftment of edited cells, as reducing AAV6 is known to reduce p21 response in HSPCs.^63^ Two donors of CB CD34^+^ cells were edited for knock-in of both 10-1074 and Ibalizumab and transplanted into non-conditioned NBSGW immunodeficient mice. Input cells carried a knock-in frequency averaging 50% with efficient knock-in of both 10-1074 and Ibalizumab constructs (Fig. 4a-b). Human chimerism and lineage formation were analyzed in the bone marrow at 12 weeks post-transplantation. Gene-edited cells engrafted at a lower frequency than mock edited cells, though this decrease is in line with those seen in the literature (Fig. 4c).^50,59–62^ As expected when human chimerism is low, gene-edited HSPCs showed a myeloid bias in reconstitution within the bone marrow (Fig. 4d).^50^ Importantly, we found the allelic knock-in frequency in the endpoint engrafted cells to remain above 50% (52-80%) in all but one mouse (11%) (Fig. 4e). Knock-in frequency was also maintained in the CD19^+^ B cell lineage population within the bone marrow, with consistent frequencies of both 10-1074 and Ibalizumab constructs, further demonstrating that the knock-in of our constructs does not impact early B cell formation (Fig. 4e and Supplementary Fig. 4b-c). Finally, *CCR5* KO INDELs were also maintained in non-integrated alleles (Fig. 4f). These results demonstrate that AZD7648 can be used to increase the frequency of edited alleles within the long-term engrafted cell population. Such an improvement provides a favorable editing profile for introducing both *CCR5* KO and long-term expression from our therapeutic antibody cassettes.

**Figure 4:**
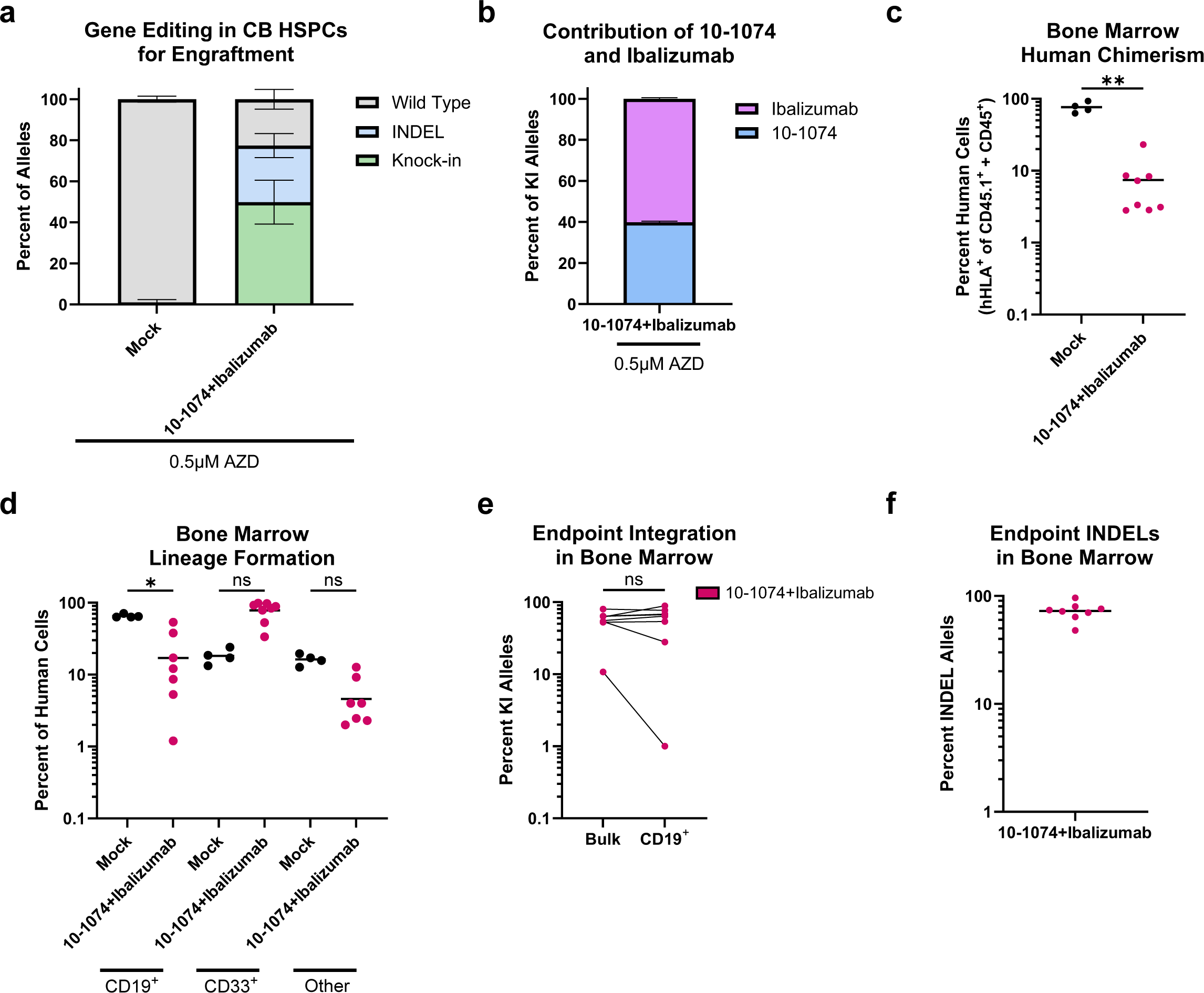
HSPCs with high frequency knock-in maintain edited alleles following engraftment *in vivo*. **a)** Distribution frequency of WT, INDEL, and KI alleles in CB CD34^+^ HSPCs prior to transplantation in NBSGW mice. Cells were targeted at *CCR5* with an AAV6 MOI of 625 for each linker antibody construct and treated with 0.5 µM AZD7648 (n = 2 independent biologic donors; data also shown in Fig. 2f). **b)** Percent of KI alleles integrated with 10-1074 or Ibalizumab within the 10-1074+Ibalizumab targeted cells shown in panel a (n = 2). **c)** Percent human cell chimerism in the bone marrow 12 weeks post-transplantation, Two-tailed Mann-Whitney test (**P = 0.0040, n = 4 for mock and n = 8 for 10-1074+Ibalizumab). **d)** Percent of human cells in the bone marrow that are CD19^+^ (B cell lineage), CD33^+^ (myeloid cell lineage), or within other lineages in mice engrafted with mock (black) or gene edited (red) HSPCs (n are the same as in c). One-way ANOVA Kruskal-Wallis test plus Dunn’s multiple comparisons test (ns, not significant, P = 0.1650 for CD33+ comparison, P = 0.3492 for Other comparison; *P = 0.0474). **e)** Percent of human alleles with knock-in from the bulk bone marrow or in positively selected bone marrow CD19^+^ cells. Lines connect dots representing measurements from the same mice (n are the same as in c). Two-tailed Mann-Whitney test (ns, not significant, P = 0.7178). **f)** Percent of human alleles from the bone marrow with an INDEL at *CCR5* (n are the same as in c). This analysis does not include alleles with KI. All bars represent mean and all error bars represent SD. All dots represent individual mice engrafted with mock HSPCs or HSPCs edited with AAV6 and RNP.

While we were able to achieve efficient engraftment of gene-edited cells, this model limits our ability to effectively analyze the *in vivo* capacity of our system to secrete antibodies. It is well known that human HSPCs transplanted into NSG background mice do not efficiently mature into highly differentiated human B cells.^64,65^ This is highlighted by our results and others showing that even mice engrafted with high human chimerism do not produce relevant levels of human IgG (Supplementary Fig. 5).^65–67^ While both humans and immunocompetent mice are expected to produce over 6 mg/mL of IgG, we observed less than 300ng/mL (>4 logs lower) in the serum of humanized mice in all but one mouse (4.5 µg/mL, 3 logs lower). Therefore, other models beyond xenotransplantation in immunodeficient mice are needed to assess the functional secretion of antibodies within our system.

### Engineered Human B cells secrete functional levels of each HIV-1 inhibiting antibody

Due to limitations in the humanized mouse model, we were unable to directly demonstrate if B cells derived from engineered HSPCs produce therapeutically relevant concentrations of linker antibodies *in vivo*. To bridge this gap, we sought to directly edit adult peripheral blood CD19^+^ B cells at the *CCR5* locus with the same RNP/AAV6 guide RNA and antibody constructs used in HSPCs. We first confirmed the function of the EEK promoter in adult B cells through knock-in of constructs driving GFP expression through either the EEK promoter or the strong ubiquitous ubiquitin C (UBC) promoter. Flow cytometry analysis for median fluorescent intensity (MFI) of GFP^+^ cells was higher when driven by the EEK promoter, demonstrating its strong activity in B cells (Supplementary Fig. 6). Next, we edited B cells with each linker antibody construct individually or with the 10-1074 and Ibalizumab constructs together. As with CD34^+^ HSPCs, we achieved highly efficient knock-in into the *CCR5* locus at frequencies up to 43% of alleles (Fig. 5a), with 10-1074 and Ibalizumab constructs integrating at a similar frequency when used in combination (Fig. 5b). Additionally, the knock-in of an exogenous antibody cassette did not obviously impact any B cell subset within the *in vitro* culture system (Supplementary Fig. 7).

**Figure 5:**
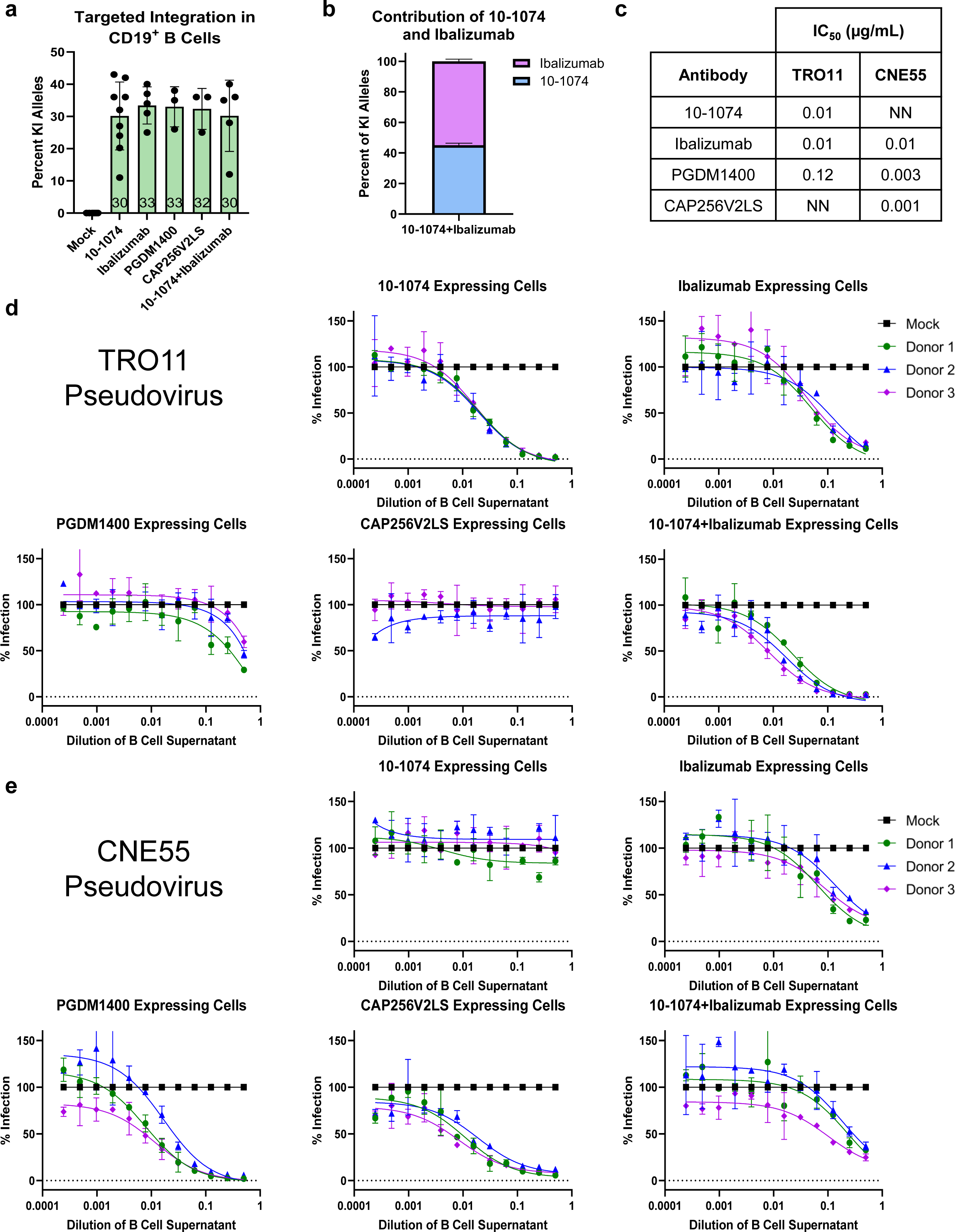
Antibody engineered B cells secrete functional linker antibodies. **a)** Allelic integration frequency in adult peripheral blood CD19^+^ B Cells targeted with AAV6 cassettes for linker antibodies as indicated. Each AAV6 was used at an MOI of 25,000 (dots represent independent biologic donors, n = 9 for mock and 10-1074, n = 5 for Ibalizumab and 10-1074+Ibalizumab, n = 3 for PGDM1400 and CAP256V2LS). **b)** Percent of KI alleles integrated with 10-1074 or Ibalizumab within the “10-1074+Ibalizumab” targeted cells shown in panel a (n = 5). **c)** Measured IC_50_ for each antibody against TRO11 or CNE55 pseudoviruses as determined by TZM-bl assay in Fig. 1c (NN, not-neutralizing). **d)** Inhibition of infection with TRO11 or **e)** CNE55 HIV-1 pseudovirus by culture supernatant from B cells engineered to express linker antibodies as indicated. Percent infection for each dose of supernatant from gene targeted B cells is normalized to infection at each dose of supernatant from mock B cells from the same donor (n = 3 biological donors, data points and error bars represent mean with standard deviation of technical duplicate infections).

We next demonstrated that the knock-in of the EEK-driven linker antibody constructs resulted in the secretion of functional antibodies. B cells were edited, allowed to recover for six days, and then plated at a density of 1 million cells per mL of media for five days. Culture supernatant was collected, and antigen-specific ELISA analysis was used to confirm the presence of each antibody (Supplementary Fig. 8). Serial dilutions of cell culture supernatant were then tested for neutralization potency in the TZM-bl infection assay against either TRO11 or CNE55 HIV-1 pseudoviruses for which we empirically determined the expected IC_50_ for each antibody in Fig. 1 (Fig. 5c). Supernatant from B cells edited with 10-1074 and Ibalizumab cassettes individually inhibited infection, with near complete inhibition at a 1:2 media dilution (Fig. 5d). As expected from the known IC_50_ values, PGDM1400 supernatant showed a reduced inhibition capacity and CAP256V2LS supernatant did not inhibit infection with TRO11. Supernatant from B cells expressing both 10-1074 and Ibalizumab effectively inhibited TRO11 infection with a similar potency to that of cells edited for each antibody alone. When tested against CNE55, we found that supernatant from B cells expressing 10-1074 did not inhibit infection, while supernatant from B cells expressing Ibalizumab, PGDM1400, and CAP256V2LS each effectively inhibited infection corresponding with the known IC_50_ of each antibody (Fig. 5e). Importantly, while cells edited to express 10-1074 alone failed to inhibit infection, cells edited to express both 10-1074 and Ibalizumab together effectively inhibited CNE55, highlighting the importance of employing multiple antibodies to overcome deficiencies of any single antibody. These results demonstrate that B cells carrying linker antibody expression cassettes are capable of effectively secreting inhibitory concentrations of antibodies individually and in combination, thus highlighting the significance of a multi-epitope targeting strategy for efficacy against diverse HIV-1 strains.

## Discussion

The transplantation of HSPCs that generate HIV-resistant progeny represents a promising strategy to achieve a functional cure for HIV-1. However, the currently established method for allogeneic transplantation of *CCR5* KO HSPCs is limited due to the rarity of available donor cells, the risk for GvHD, and a lack of efficacy against X4-tropic virus. Transplantation of genetically engineered autologous HSPCs is a treatment modality with the potential to address these limitations. Modifying a patient’s own cells removes the need for identifying a matched donor and circumvents the risks specific to allogeneic transplantation. Additionally, precise genetic engineering can allow for the delivery of multi-factored resistance to R5-tropic and X4-tropic HIV-1. To this end, we have established a highly efficient genome editing strategy to combine the known benefits of *CCR5* KO with antibody therapy in a manner applicable towards autologous HSPC transplantation.

Here we demonstrate the simultaneous knockout of *CCR5* and knock-in of antibody expression cassettes in human HSPCs for secretion of HIV-1 inhibiting antibodies in B cell progeny of edited cells. Following transplantation, genetically engineered HSPCs have the potential to persist for a patient’s lifetime and provide a long-term source of therapeutic antibody secreting B cells. Recent trials investigating direct injection of bNAbs for HIV-1 suppression show that repeated dosing is required to maintain efficacy, as viral titers rebound once the serum concentrations drop below a therapeutic threshold.^18,19^ Additionally, AAV and B cell-mediated antibody delivery platforms rely on somatic cells and are unlikely to rival the longevity of HSCT. Therefore, the engraftment of genetically engineered HSPCs represents one of the most promising methods for a one-time therapy to combat HIV-1. To this end, we show that RNP/AAV6 editing of HSPCs delivers a favorable profile of modified alleles, achieving a knock-in frequency of up to 50% of alleles and KO frequency of up to 90% of alleles at *CCR5* (Fig. 2). Importantly, engineered HSPCs maintain their ability to engraft and reconstitute the hematopoietic lineages in immunodeficient mice (Fig. 3), and small molecule inhibition of NHEJ can be used to address the decrease in knock-in alleles normally seen following engraftment (Fig. 4). The development of this system for highly efficient *CCR5* KO and knock-in of antibody expression cassettes is a favorable first step towards developing an autologous transplantation therapy.

Recently, the transplantation of gene edited autologous HSPCs has displayed clinical success with the approval of Casgevy for the treatment of sickle cell disease and beta-thalassemia.^68–70^ Casgevy uses RNP editing for INDEL generation in HSPCs, acting as a proof of concept for RNP-edited HSCT. Now, efforts are underway to expand the therapeutic potential of CRISPR-Cas9 editing to include knock-in of large therapeutic cassettes using AAV6/RNP mediated editing. However, it is regularly shown that AAV6 treatment negatively impacts HSPCs, as demonstrated through reduced colony formation in CFU assays and decreased chimerism following xenotransplantation.^71^ We and others are exploring methods to alleviate toxicity associated with AAV6 in HSPCs to improve the long-term engraftment of engineered cells. For one, the use of small molecule NHEJ inhibitors allows for reduced doses of AAV6 and RNP to achieve the same editing frequencies, potentially decreasing toxicity through a reduction in p53 activation.^58,63,72^ Additionally, inhibition of the cell’s DNA damage response through transient inhibition of 53BP1 with peptides or siRNA has been shown to reduce toxicity in HSPCs.^72,73^ Finally, it has been demonstrated that simply increasing the number of transplanted cells can compensate for AAV6 associated toxicity in xenograft mouse models.^61^ While more work is needed to refine optimal editing conditions and improve engraftment of AAV6/RNP-edited HSPCs, the data presented here represent an important proof of concept study for the efficacy of a long-term antibody delivery platform for treatment of HIV and other diseases treatable by recombinant antibody delivery.

Engraftment in immunodeficient mice is the gold-standard for functional analysis of genetically engineered human HSPCs, however, the model fails to recapitulate normal B cell development and maturation. While our engineered HSPCs do engraft and maintain edited alleles in CD19^+^ progeny, we are not able to assess functional antibody expression due to a lack of mature antibody secreting B cells *in vivo*. To address our system’s ability to drive therapeutic antibody expression, we edited adult peripheral blood B cells with the same constructs used in HSPCs. These modified B cells were capable of secreting inhibiting levels of each tested antibody into the culture supernatant (Fig. 5). It is well known that multiple antibodies should be employed simultaneously to combat viral diversity and escape. We therefore designed our system for use with multiple antibodies concurrently and show that linker 10-1074 and linker Ibalizumab could be expressed together from one bulk population of B cells (Fig. 5 and Supplementary Fig. 8). The modular design of our cassettes permits antibodies to be readily switched out, allowing for optimization of antibody pairings and potential customization based on the viral diversity of various patient populations across regions.

Because B cells are the professional antibody-producing cells of the hematopoietic system, we believe that expression from B cell progeny is a favorable strategy for sustained delivery of engineered antibodies. By expressing from the *CCR5* locus, our cassettes will act as passengers to normal B cell function and should therefore persist across B cell sub-types. This will ideally allow for our antibodies to reach a steady-state concentration in the bloodstream, providing an advantage over traditional antibody injection therapies that have peaks and troughs with each dosage. Additionally, expression of these antibodies from B cells directly may benefit from the inclusion of natural post-translational modifications that are not incorporated when expressed from other cell types.^74^ Carrying these modifications may decrease immune recognition of HIV-1 inhibiting antibodies and reduce the potential for anti-drug antibody formation, an issue encountered by both vectored immunoprophylaxis and direct injection trials.^24,27,75,76^ Additionally, endogenous glycosylation and sialylation have the potential to improve F_C_-mediated clearance of HIV-1 infected cells.^77,78^ Moving forward, it will be important to investigate the impact of post-translational modifications on antibodies expressed from B cells.

Overall, this work serves as a platform strategy for the lifetime secretion of desired therapeutic antibodies. Beyond its usefulness for HIV-1 resistance, knock-in into the *CCR5* safe harbor site allows for broad applications across chronic diseases. Future works will test alternate antibodies for use within our system as a new delivery modality for current treatment regimens that require long-term dosing of monoclonal antibodies.

## Methods

### rAAV6 vector design, production, and purification

The sequence of each antibody construct was cloned from a gblock Gene Fragment (Integrated DNA Technologies, IDT, San Jose, CA, USA). Restriction enzyme digest or Gibson Assembly (New England Biolabs, Ipswich, MA, USA) was used to clone each antibody sequence plus 400 base pair (bp) homology arms on each side of the construct into an adeno-associated virus, serotype 6 (AAV6) vector plasmid derived from the pAAV-MCS plasmid (Agilent Technologies, Santa Clara, CA, USA). Experiments were performed with rAAV6 vectors produced and purified by SignaGen Laboratories (Frederick, MD, USA) and Packgene Biotech (Houston, TX, USA). All viral titers were determined using droplet digital PCR (ddPCR) to measure the number of vector genomes as previously described.^79^

### CD34^+^ HSPC isolation and culture

Human CD34^+^ HSPCs were isolated from cord blood by the Stanford Binns Program for Cord Blood Research and cultured as previously described.^80^ Briefly, isolated mononuclear cells were positively selected for CD34 using the CD34+ Microbead kit Ultrapure (Miltenyi Biotec, San Diego, CA, USA, cat.: 130-100-453). Cells were cultured at 1.5×10^5^–2.5×10^5^ cells/mL in CellGenix® GMP Stem Cell Growth Medium (SCGM, CellGenix, Freiburg, Germany, cat.: 20802-0500) supplemented with a human cytokine (PeproTech, Rocky Hill, NJ, USA) cocktail: stem cell factor (100 ng/mL), thrombopoietin (100 ng/mL), Fms-like tyrosine kinase 3 ligand (100 ng/mL), interleukin 6 (100 ng/mL), streptomycin (20 mg/mL), and penicillin (20 U/mL), and 35 nM of UM171 (APExBIO, Houston, TX, USA, cat.: A89505). Cells were cultured in a 37°C hypoxic incubator with 5% CO_2_ and 5% O_2_. Cells were cultured for 3 days prior to editing.

### Genome editing of HSPCs

HPLC-purified synthetic chemically modified sgRNAs were purchased from TriLink Biotechnologies (San Diego, CA, USA). Chemical modifications were comprised of 2′-O-methyl-3′-phosphorothioate at the three terminal nucleotides of the 5′ end and the second, third, and fourth bases from the 3’ end as described previously.^81^ The target sequence for the sgRNA are as follows: *sg-CCR5,* 5′-ATGCACAGGGTGGAACAAGA-3′; *sg-CCR5-#2*, 5’-GCAGCATAGTGAGCCCAGAA-3’. HiFi Cas9 protein was purchased from IDT (cat.: 1081061) or Aldevron (Fargo, ND, USA, cat:. 9214). RNPs were complexed at a Cas9:sgRNA molar ratio of 1:2.5 at room temperature for 15-30 min. 2.5×10^5^-1×10^6^ CD34^+^ cells were resuspended in P3 buffer (Lonza, Basel, Switzerland, cat.: V4XP-3032) with complexed RNPs and electroporated using the Lonza 4D Nucleofector and 4D-Nucleofector X Unit (program DZ-100). Electroporated cells were then plated at 2.5×10^5^ cells/mL in the previously described cytokine-supplemented media. Immediately following electroporation, AAV6 was dispensed onto cells at 0.625×10^3^-2.5×10^3^ vector genomes/cell as noted in figure legends. For some editing experiments, in addition to the steps described above, cells were incubated with 0.5 μM of DNA-PKcs inhibitor, AZD7648 (Selleck Chemicals, Houston, TX, cat.: S8843) for 24 hours, as previously described.^58^

### B cell isolation, culture, and genome editing

Leukoreduction system (LRS) chambers were obtained from the Stanford Blood Center and primary human B cells were isolated by negative selection using the human B Cell Isolation Kit II (Miltenyi Biotec, cat: 130-091-151). Cells were cultured in Iscove’s modified Dulbecco’s medium (IMDM) (Thermo Fisher Scientific, Waltham, MA, USA cat.: 12440053) supplemented with 10% bovine growth serum (Cytiva, Marlborough, MA, USA, cat.: SH30541.03HI), 1% penicillin-streptomycin (Cytiva, cat.: SV30010), 55 µM of 2-mercaptoethanol (Sigma-Aldrich, St. Louis, MO, USA, cat.: M3148), 50 ng/mL of IL-2 (Peprotech, cat.: 200-02), 50 ng/mL of IL-10 (Peprotech, cat.: 200-10), 10 ng/mL of IL-15 (Peprotech, cat.: 200-15), 100 ng/mL of recombinant human MEGACD40L (Enzo Life Sciences, Farmingdale, NY, USA, cat.: ALX-522-110-C010), and 1 µg/mL of CpG oligonucleotide 2006 (Invivogen, San Diego, CA, USA cat.: tlrl-2006-1) at a density of 1×10^6^ cells/mL, as described previously.^28^ B cells were cultured at 37°C, 5% CO_2_, and ambient oxygen levels. Genome editing was performed as described above for HSPCs with the following modifications. B cells were edited 3-5 days after isolation or thawing using the Lonza Nucleofector 4D (program EO-117) using 1×10^6^ cells per well of a 16-well Nucleocuvette Strip (Lonza). Immediately following nucleofection, cells were incubated with AAV6 donor vector (2.5×10^4^ vector genomes/cell) in 100 µl of basal IMDM in a 96 well plate for 3-4 hours and then replated at 1×10^6^ cells/mL in complete B cell activation media.^82^

### Measurement of knock-in alleles by ddPCR

Cells were harvested 2-3 days post-electroporation and genomic DNA (gDNA) was harvested using QuickExtract DNA extraction solution (Biosearch Technologies, Hoddesdon, UK, cat.: QE09050). To quantify knock-in alleles via ddPCR, we employed *CCR5* specific in-out PCR primers and a probe corresponding to the expected knock-in event (1:3.6 primer to probe ratio).^60^ We also used an established genomic DNA reference (REF) at the *CCRL2* locus.^60^ The ddPCR reaction was prepared and underwent droplet generation following the manufacturer’s instructions with a Bio-Rad QX200 ddPCR machine (Bio-Rad, Hercules, CA, USA). Thermocycler settings were as follows: 95°C (10 min, 1°C/s ramp), 94°C (30 s, 1°C/s ramp), 60°C (30 s, 1°C/s ramp), 72°C (2 min, 1°C/s ramp), return to step 2 for 50 cycles, and 98°C (10 min, 1°C/s ramp). Analysis of droplet samples was then performed using the QX200 Droplet Digital PCR System (Bio-Rad). We next divided the copies/μL for HDR (%): HDR (FAM) / REF (HEX). The following primers and probes were used in the ddPCR reaction:

*CCR5-BGH:*

Forward Primer (FP): 5’-GGGAGGATTGGGAAGACA-3’

Reverse Primer (RP): 5’-AGGTGTTCAGGAGAAGGACA-3’

Probe: 5’-6-FAM/AGCAGGCATGCTGGGGATGCGGTGG/3IABkFQ-3’

*CCR5-IgG1:*

FP: 5’-CCTGAGCCCCGGAAAATAG-3’

RP: 5’-AGGTGTTCAGGAGAAGGACA-3’

Probe: 5’-6-FAM/AGCAGGCATGCTGGGGATGCGGTGG/3IABkFQ-3’

*CCR5-IgG4:*

FP: 5’-CCTCTCCCTGTCTCTGGGTA-3’

RP: 5’-AGGTGTTCAGGAGAAGGACA-3’

Probe: 5’-6-FAM/AGCAGGCATGCTGGGGATGCGGTGG/3IABkFQ-3’

*CCRL2:*

FP: 5’-GCTGTATGAATCCAGGTCC-3’,

RP: 5’-CCTCCTGGCTGAGAAAAAG-3’

Probe: 5’-HEX/TGTTTCCTC/ZEN/CAGGATAAGGCAGCTGT/3IABkFQ-3’

### INDEL analysis using ICE software

Two-or three-days following editing, cells were collected and genomic DNA was extracted using QuickExtract DNA extraction solution (Biosearch Technologies, cat.: QE09050). The following primer sequences were used to amplify the *CCR5* locus around the sgRNA cut site as previously described:^50^

FP: 5’-CAGGGAAGCTAGCAGCAAACC-3’

RP: 5’-AGACGCAAACACAGCCACC-3’

PCR products were run on a 1% agarose gel and purified using the GeneJET Gel Extraction Kit (Thermo Fisher Scientific, cat.: FERK0692). Sanger sequencing of respective PCR samples was performed using the reverse primer. Sequencing files were then used as input for INDEL frequency analysis relative to a mock, unedited sample using the online ICE CRISPR analysis tool (Synthego, Redwood City, CA, USA).^83^

### PCR and Gel Electrophoresis for Knock-in

Two-or three-days following editing, cells were collected and genomic DNA was extracted using QuickExtract DNA extraction solution (Biosearch Technologies, cat.: QE09050). In-out PCR was performed with 3 forward primers and one reverse primer to selectively amplify knock-in events of the Ibalizumab, 10-1074, and PGDM1400 antibody cassettes. Primers used were as follows:

Ibalizumab FP: 5’-ACAGTCCTCAGGACTCTACTCC-3’

10-1074 FP: 5’-TATGGCGTGGTGAGCTTTGG-3’

PGDM1400 FP: 5’-CTGGGACCTCCGTAAAGGTCT-3’

CCR5 RP: 5’-AGACGCAAACACAGCCACC-3’

PCR products were run on a 1% agarose gel and imaged using a Bio-Rad ChemiDoc XRS+ gel imager.

### Antibody production and purification

Full length heavy and light chains for each antibody were cloned by restriction enzyme digest or Gibson Assembly (New England Biolabs) into pCMVR either individually or with a linker. Antibody constructs were expressed in Expi293F cells (Thermo Fisher Scientific, cat.: A14527) and grown in combined media (66% FreeStyle/33% Expi media, Thermo Fisher Scientific, cat.: 12338018 and A1435101, respectively) at 37°C and 8% CO2 with shaking at 120 rpm. Antibody constructs were transfected using FectoPRO transfection reagent (Polyplus, Sébastien Brant, France, cat.: 101000014) with a transfection mixture of 10 mL of culture media, 50 µg of total DNA and 130 µL of FectoPro for each 90 mL of cell culture (for a total transfected cell culture volume of 100 mL or scaled down at the same ratios for a 75 mL total volume). Heavy and light chain plasmids were used at a 1:1 ratio, while single plasmid linker antibody constructs were used alone. Cells were transfected at a cell density 3-4×10^6^ cells/mL and harvested 4 days post-transfection by spinning at >4200g for 15 min. Cell culture supernatants were filtered through a 0.45-μm filter, combined with 1/10^th^ volume of 10x phosphate-buffered saline (PBS) and purified using the ÄKTA pure fast performance liquid chromotography (FPLC, Cytiva) with a 5mL MabSelect PrismA column (Cytvia, cat.: 17549802) using wash steps with 1x PBS and elution with 100 mM glycine (pH 2.8) into one-tenth volume of 1 M Tris (pH 8.0). The eluted proteins were then concentrated using 50-kDa or 100-kDa cutoff centrifugal concentrators and further purified by size exclusion on the same ÄKTA FPLC with Superdex 200 Increase 10/300 GL column (Cytiva, cat.: 28-9909-44). Proteins were then further concentrated using 50-kDa or 100-kDa cutoff centrifugal concentrators.

### Immunophenotyping of B cells

All samples were blocked for non-specific binding with human FcR Blocking Reagent (Miltenyi Biotec, cat.: 130-059-901) for 10 min at room temperature and then stained for 30 min at 4°C with the following antibody cocktail: anti-human CD19 (Becton, Dickinson and Company, BD, Franklin Lakes, NJ, USA, cat.: 562440), anti-human CD20 (BioLegend, San Diego, CA, USA, cat.: 302326), anti-human CD27 (BioLegend, cat.: 302832), anti-human CD38 (BD, cat.: 555462), anti-human IgD (BioLegend, cat.: 348232), and anti-human IgM (BD, cat.: 563903). Samples were analyzed on a FACSAria II SORP (BD). Data was subsequently analyzed using FlowJo (v.10.9.0).

### ELISA for Antibody Concentration

Antigen specific ELISA was used to determine the concentration of each HIV-inhibiting antibody in B cell supernatant. Anti-IgG Fc ELISA was used to detect human IgG in mouse serum. 10-1074, Ibalizumab, and total human IgG were detected using a previously described protocol.^84^ Briefly, Nunc MaxiSorp 96 Well Plates (Thermo Fisher Scientific, cat.: 44-2404-21) were coated with 0.8 µg/mL recombinant HIV_JR-CSF_ gp120 (for detecting 10-1074; Immune Technology, Tarrytown, NY, USA, cat.: IT-001-0025p), 1 μg/mL recombinant rhesus monkey CD4 protein (for detecting Ibalizumab; Abcam, Cambridge, UK, cat.: ab208305), or 1:100 diluted goat anti-human IgG-Fc (for detecting total human IgG; Fortis Life Sciences, Waltham, MA, USA, cat.: A80-104A) in 0.05 M carbonate-bicarbonate buffer solution (Sigma-Aldrich, cat.: C3041) for 1 hour at room temperature or overnight at 4°C. Plates were then blocked in tris-buffered saline (TBS) with 1% bovine serum albumin (BSA; Miltenyi Biotech, cat.: 130-091-376) for 30 minutes. Media and serum samples were diluted in TBS with 1% BSA and 0.05% Tween-20 and incubated on the plate for 4 hours at room temperature or overnight at 4°C. Plates were then incubated with horseradish peroxidase (HRP)–conjugated goat anti-human IgG-Fc antibody (Fortis Life Sciences, cat.: A80-104A) at a 1:2500 dilution.

Detection was performed with the 1-Step™ TMB Substrate Kit (Thermo Fisher Scientific, cat.: 34021) and quenched with 3M H_2_SO_4_. Plates were read at 450nm using a Molecular Devices SpectraMax M3 plate reader with SoftMax Pro software. Standard curves were created using purified versions of each HIV inhibiting antibody or purified human IgG/kappa from normal serum (Bethyl, Montgomery, TX, USA, cat.: P80-111). Results were analyzed using GraphPad Prism v10 software to calculate a standard curve using a 4-parameter or 5-parameter sigmoidal algorithm. The curve with a higher r-squared was selected and results were interpolated for each sample. PGDM1400 and CAP256V2LS were detected with a modified protocol. Briefly, recombinant HIV-1 Env trimer, BG505 SOSIP, was biotinylated with the EZ-Link™-Biotinylation Kit (Thermo Fisher Scientific, cat.: 21435) as previously described.^85^ Nunc MaxiSorp plates were coated with 4 µg/mL streptavidin (Thermo Fisher Scientific, cat.: PI21122) in PBS at room temperature for 1 hour. Plates were then blocked with ChonBlock (Thermo Fisher Scientific, cat.: 50-152-6971) overnight at 4°C. The next day, 1 µg/mL biotinylated BG505 SOSIP trimer was added for 1 hour at room temperature. Samples were diluted in PBS with 0.1% BSA and 0.05% Tween-20 and incubated on the coated plates for 1 hour. Plates were then detected with HRP-conjugated anti-human IgG-Fc antibody as described above. Results were analyzed as described above.

### Western Blotting of Purified Antibodies

Purified antibodies (0.5 ug each) were denatured by adding 4X LaemmLi sample buffer (Bio-Rad, cat.: 1610747) and 2-mercaptoethanol and heating at 100°C for 5 minutes. Samples were loaded onto a 4-15*%* Mini-PROTEAN TGX precast protein gel (Bio-Rad, cat.: 4561084). Following electrophoresis, protein from gel was transferred to a PVDF membrane using the Trans-Blot Turbo Transfer System (Bio-Rad, cat.: 1704150).

Subsequently, the membrane was blocked using Blotting-Grade Blocker (Bio-Rad, cat.: 1706404) in phosphate-buffered saline with 0.02% Tween 20 (PBS-T) for 30 minutes at room temperature. Membrane was then incubated in 1:5000 primary antibody (HRP-conjugated Goat pAb to human IgG, Abcam, cat.: 7153) overnight at 4°C. The next day, the membrane was washed three times with PBS-T. SuperSignal™ West Pico PLUS Chemiluminescent Substrate (Thermo Fisher Scientific, cat.: 34579) was used for detection and blots were imaged using a Bio-Rad ChemiDoc XRS+ gel imager using ImageLab software.

### TZM-bl Infection Assay for Determining IC_50_

The TZM-bl HIV-1 pseudotype infection assay was adapted from previously described protocols.^86,87^ TZM-bl cells were obtained through the NIH AIDS Reagent Program (cat.: 8129) and cultured in DMEM with 10% BGS, and 1% penicillin-streptomycin (complete DMEM) at 37°C, 5% CO2, and ambient oxygen levels. HIV-1 env pseudotyped lentivirus was produced as previously described using a pNL4-3ΔEnvGFP backbone plasmid and *env* plasmids provided by the NIH AIDS Reagents Program (cat.: 11100 and cat.: 12670, respectively).^88^ Briefly, 5×10^3^ cells were plated in black-walled, clear-bottom 96-well plates (Corning, Corning, NY, USA, cat.: 07-200-565) and incubated overnight. The next day, each antibody was incubated with HIV-1 pseudovirus for 1 hour at 37°C at concentrations from 5 µg/mL to 5×10^−5^ µg/mL using 10-fold serial dilution in complete DMEM with 15 µg/mL DEAE-dextran (Sigma-Aldrich, D9885). Culture media was aspirated from the TZM-bl cells and replaced with antibody-virus mixture. Virus only (no antibodies added) and cells only (no virus or antibodies) wells were included for determining 100% and 0% infection readouts. Cells were incubated for 48 hours then measured for luciferase signal corresponding to infection. Briefly, cells were lysed with Reporter Lysis Buffer (Promega, Madison, WI, USA, cat.: E3971) and freeze-thawed at −80°C. Lysed samples were read for relative luminescence units (RLU) using a Synergy H1 plate reader (BioTek, Winooski, VT, USA) that automatically injected luciferin solution consisting of the following: 200 mM Tris [pH 8], 10 mM MgCl2, 300 μM ATP, Firefly Luciferase Signal Enhancer (Thermo Fischer Scientific, cat. 16180), and 150 μg/mL d-luciferin (Biosynth Chemistry & Biology, Staad, Switzerland, cat.: L8220). Percent infection was determined by normalizing RLU values to the average RLU of virus-only and cells-only wells using GraphPad Prism v10. IC_50_ was calculated using a linear regression dose-response curve fit for inhibitor versus response (three parameters) based on the average of technical duplicate wells using GraphPad Prism v10.

### TZM-bl Infection Assay with B cell supernatant

Six days post-editing, B cells were plated at 1×10^6^ cells/mL and antibody was allowed to accumulate in supernatant for five days. Culture supernatant was collected following centrifugation to remove cells. A modified TZM-bl assay was performed as described above with the following changes. Culture supernatant was diluted in complete DMEM with two-fold serial dilutions. Then, 50uL of each dilution was mixed with 50uL of HIV-1 pseudovirus + complete DMEM for final dilutions of culture media ranging from 1:2 to 1:4096 and a final concentration of 15 µg/mL DEAE-dextran (Sigma-Aldrich, cat.: D9885). Samples were incubated for 1 hour at 37°C before being applied to TZM-bl cells.

### Colony forming unit (CFU) assay and colony genotyping

Two days post-electroporation 500 HSPCs were plated in each well of a SmartDish 6-well plate (STEMCELL Technologies, Vancouver, Canada, cat.: 27370) containing MethoCult H4434 Classic (STEMCELL Technologies, cat.: 04444). After 14 days, the wells were imaged using the STEMvision Hematopoietic Colony Counter (STEMCELL Technologies). Colonies were counted and scored with manual correction to determine the number of BFU-E, CFU-GM, and CFU-GEMM colonies. Individual colonies were picked and gDNA was extracted using QuickExtract DNA extraction solution (Biosearch Technologies, cat.: QE09050). Knock-in was determined by ddPCR and INDELs were determined by ICE analysis as described above.

### Mice

NOD.Cg-Prkdc^scid^Il2rg^tm1Wjl^/SzJ (NSG) and NOD.Cg-*Kit^W-41J^Tyr*^+^*Prkdc^scid^ Il2rg^tm1Wjl^*/ThomJ (NBSGW) mice were purchased from Jackson Laboratories (Bar Harbor, ME, USA). Mice were housed in the Stanford University barrier facility. All experiments were completed under the Administrative Panel on Laboratory Animal Care (APLAC Protocol #25065).

### CD34^+^ HSPC transplantation in to immunodeficient mice

For the experiment in NSG mice, 6-8 week old mice were irradiated with 2Gy approximately 4 hours prior to transplantation. 8.5×10^5^ mock or gene-edited cells were transplanted into each mouse via retro-orbital injection. For the experiment in NSGBW mice, 6-8 week old mice received no conditioning and 5×10^5^ mock or gene-edited cells were transplanted into each mouse via retro-orbital injection.

### Assessment of human HSPC engraftment

Human engraftment was assessed at 12-(NSGBW) or 16-weeks (NSG) post-transplantation. Mice were euthanized and bone marrow and spleen were harvested from recipient mice. Mononuclear cells from bone marrow samples were isolated via Ficoll gradient centrifugation. Spleen samples were treated with RBC Lysis Buffer (IBI Scientific, Dubuque, IA, USA, cat.: IB47620) to eliminate mature red blood cells. All samples were blocked for non-specific binding with FcR Blocking Reagent for human (Miltenyi Biotec, cat.: 130-059-901) and mouse (Miltenyi Biotec, cat.: 130-092-575) for 10 min at room temperature and then stained for 30 min at 4°C with the following antibody cocktail: anti-mouse CD45.1 (Biolegend, cat.: 110735), anti-mouse TER-119 (eBiosciences, San Diego, CA, USA, cat.: 15-5921-82), anti-human CD45 (Biolegend, cat.: 368540), anti-human HLA-ABC (Biolegend, cat. 311426), anti-human CD33 (BD, cat.: 555450), anti-human CD19 (Biolegend, cat.: 302212), anti-human CD3 (Biolegend, cat.: 300328), and viability dye Ghost Dye™ Violet 540 (Tonbo Bioscience, San Diego, CA, USA, cat. 130879T100). Samples were analyzed on a CytoFLEX flow cytometer (Beckman-Coulter, Indianapolis, IN, USA). Data was subsequently analyzed using FlowJo (v.10.9.0). Humam CD19^+^ cells were isolated from the bone marrow using positive selection with the CD19 MicroBeads, human kit (Miltenyi Biotec, cat.: 130-050-301). Knock-in and INDEL frequencies were determined by ddPCR and ICE analysis as described above.

### Statistical analysis

All statistical analysis was performed using GraphPad Prism v10 software.

## Data availability

Reagents and protocols are available upon request from the corresponding author. Source data will be provided with the completed manuscript.

## Acknowledgements

We thank Dr. Peter Kim for technical advice and support in purifying and testing antibodies. We thank Dr. Christopher Barnes for technical advice and for providing the BG505 SOSIP Trimer. We thank Dr. Alejandro Balazs for assistance with the gp120 ELISA. We thank the NIH AIDS Reagent Repository for providing the *env* plasmids for the panel of HIV-1 pseudoviruses. We thank the Stanford Institute for Stem Cell Biology and Regenerative Medicine FACS Core for their support and access to the flow cytometry machines. We thank the Binns Program for Cord Blood Research for providing purified cord blood HSPCs. W.N.F was supported by the Stanford T32 Graduate Training Program in Stem Cell Biology and Regenerative Medicine (5T32GM119995) and the Blavatnik Family Foundation as a Blavatnik Fellow. S.E.L was supported by the Stanford Medical Scholars Research Program, the American Society of Hematology Minority Medical Student Award Program, and the Stanford Medical Scientist Training Program. M.F.I. was supported by the Stanford Medical Scientist Training Program (5T32GM007365) and the Ruth L Kirschstein Individual Predoctoral NRSA awarded by the NIH/NIAID (5F30AI152943). T.U.J.B was supported by the Knight-Hennessy Graduate Scholarship and a Canadian Institutes of Health Research Doctoral Foreign Study Award (FRN:170770). A.M.D. was supported by NIH F32 Individual Postdoctoral Fellowship (1F32HL154667-01)) and NIH K99/R00 Pathway to Independence Award (1K99HL172253-01). M.H.P. was supported by the Sutardja Chuk Professorship in Definitive and Curative Medicine and the Laurie Kraus Lacob Translational Medicine endowment.

## Author Contributions

W.N.F., S.E.L, K.B.E, M.V.F.I, T.U.J.B, H.Y.G and A.M.D. contributed to experimental design, performance, and analysis. N.A.A. and N.M.J performed experiments for *in vivo* engraftment studies. B.J.L. contributed to the design and direction of the project. W.N.F., S.E.L, and M.H.P. wrote the draft and finalized versions of the manuscript with input from other authors. M.H.P. supervised the project.

