## Supplementary Information for "Combining Cell-Intrinsic and -Extrinsic Resistance to HIV-1 By Engineering Hematopoietic Stem Cells for CCR5 Knockout and B Cell Secretion of Therapeutic Antibodies"

**Supplementary Figure Legends:**

**Supplementary Figure 1: Inhibition activity of traditional and linker antibodies against a global panel of HIV-1 pseudoviruses. a)** Expected and measured $IC_{50}$ of each antibody. Expected $IC_{50}$ was derived from CATNAP unless otherwise noted.[89,90] Traditional and linker antibody $IC_{50}$ was determined as described in Fig. 1c (data also shown in Fig. 1c).

### Supplementary Figure 1: Inhibition activity of traditional and linker antibodies against a global panel of HIV-1 pseudoviruses

**a**

**$IC_{50}$ (μg/mL)**

| | 10-1074 | | | 3BNC117 | | | Ibalizumab | | | PGDM1400 | | | CAP256V2LS | | | 1-18 | | |
|---|---|---|---|---|---|---|---|---|---|---|---|---|---|---|---|---|---|---|
| **Pseudovirus** | **Expected** | **Original** | **Linker** | **Expected** | **Original** | **Linker** | **Expected** | **Original** | **Linker** | **Expected** | **Original** | **Linker** | **Expected** | **Original** | **Linker** | **Expected** | **Original** | **Linker** |
| **TRO11** | 0.013 | 0.010 | 0.014 | 0.046 | 0.032 | 0.496 | 0.025 | 0.016 | 0.016 | 0.503 | 0.158 | 0.120 | >5 | >5 | >5 | 0.017 | 0.022 | 0.140 |
| **25710** | 0.027 | 0.046 | 0.066 | 0.136 | 0.291 | >5 | 0.557 | 0.016 | 0.011 | 0.006 | 0.002 | 0.002 | 0.001 | 0.001 | 0.001 | 0.030 | 0.049 | 0.152 |
| **BJOX2000** | 0.013 | 0.013 | 0.015 | >5 | >5 | >5 | 0.610* | 0.015 | 0.021 | 0.002 | 0.006 | 0.010 | 0.001 | 0.002 | 0.002 | 0.040 | 0.021 | 0.086 |
| **X1632** | >5 | >5 | >5 | 3.260 | 0.290 | >5 | >5 | >5 | >5 | 0.009 | 0.014 | 0.016 | 0.001 | 0.001 | 0.001 | 0.023 | 0.028 | 0.303 |
| **CE1176** | 0.023 | 0.018 | 0.034 | 0.259 | 0.202 | >5 | 0.053 | 0.012 | 0.018 | 0.139 | 0.067 | 0.016 | 0.012 | 0.005 | 0.005 | 0.214 | 0.100 | 0.720 |
| **246F3** | >5 | >5 | >5 | 0.102 | 0.210 | 1.786 | 0.036 | 0.022 | 0.021 | 0.003 | 0.003 | 0.003 | 0.009 | 0.010 | 0.009 | 0.030 | 0.035 | 0.299 |
| **CH119** | 0.022 | 0.020 | 0.024 | 4.734 | 1.726 | >5 | 0.061 | 0.018 | 0.017 | 0.075 | 0.200 | 0.214 | 0.019 | 0.019 | 0.034 | 0.050 | 0.045 | 0.261 |
| **CE0217** | 0.008 | 0.011 | 0.009 | 0.042 | 0.068 | 0.401 | 0.244 | 0.027 | 0.027 | 0.004 | 0.007 | 0.005 | 0.003 | 0.002 | 0.003 | 0.040 | 0.039 | 0.405 |
| **CNE55** | >5 | >5 | >5 | 0.119 | 0.190 | 1.051 | 1.563 | 0.015 | 0.014 | 0.009 | 0.002 | 0.003 | 0.001 | 0.001 | 0.001 | 0.020 | 0.030 | 0.223 |

*[90]Moshoette, T., Ali, S. A., Papathanasopoulos, M. A. & Killick, M. A. Engineering and characterising a novel, highly potent bispecific antibody iMab-CAP256 that targets HIV-1. Retrovirology 16, 1-12 (2019)

| Key (μg/mL) |
|---|
| <0.01 |
| <0.1 |
| <1 |
| >1 |

**Supplementary Figure 2: *CCR5* INDEL profile and construct specific ddPCR for integration analysis. a)** Representative INDELs formed in CB CD34+ HSPCs following gene editing with *CCR5* RNP. Image generated with Synthego ICE Analysis. **b)** Diagram of ddPCR primer design for measuring bulk and construct specific integration at the *CCR5* locus. Bulk integration is based on the BGH-polyA signal that all antibody cassettes carry. Construct specific integration is based on IgG1 (10-1074) or IgG4 (Ibalizumab). **c)** Percent of KI alleles as measured by ddPCR using bulk integration or construct specific integration primers across dilutions of CB CD34+ HSPCs targeted with an IgG1 antibody construct (10-1074, n = 1). **d)** Percent of KI alleles as measured by ddPCR using bulk integration or construct specific integration primers across dilutions of CB CD34+ HSPCs targeted with an IgG4 antibody construct (Ibalizumab, n = 1). **e)** Efficiency of construct specific ddPCR primers relative to bulk integration primers. Measurement efficiency is determined by dividing the construct specific integration measurement by the bulk integration measurement at the indicated dilutions of targeted cells.

**Supplementary Figure 2: *CCR5* INDEL profile and construct specific ddPCR for integration analysis**

**a**

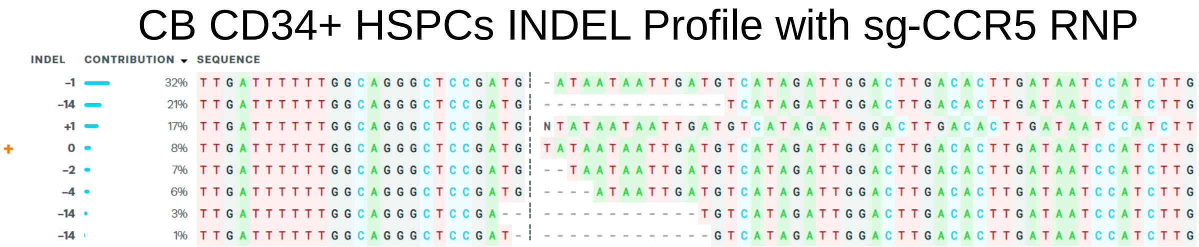


**b**

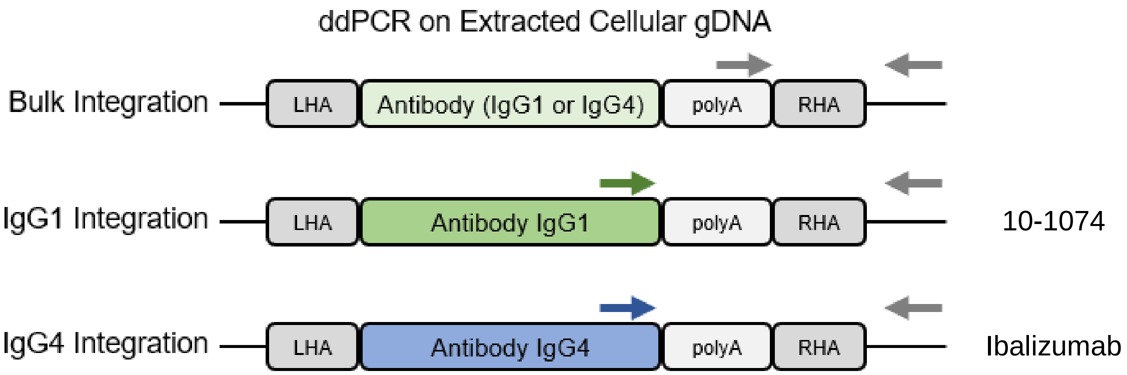


**c**

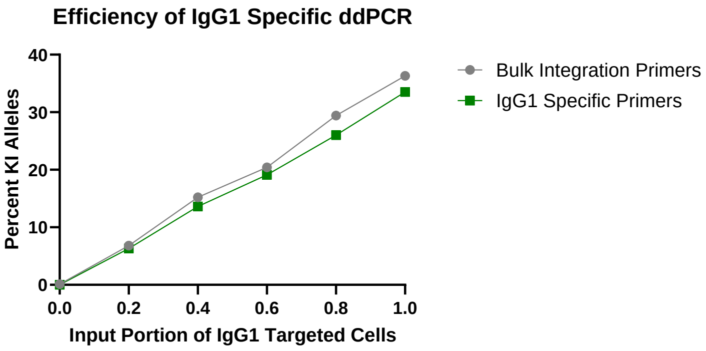


**d**

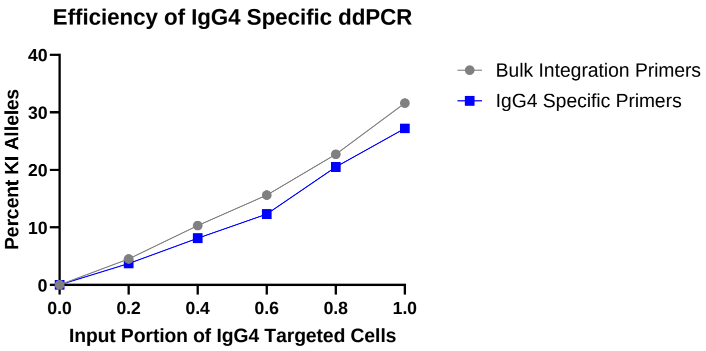


**e**

| | Input Portion of Targeted Cells | 0.2 | 0.4 | 0.6 | 0.8 | 1.0 |
|---|---|---|---|---|---|---|
| Measurement Efficiency Compared to Bulk Primers | IgG1 Specific Primers | 93% | 90% | 93% | 89% | 92% |
| | IgG4 Specific Primers | 83% | 79% | 79% | 90% | 86% |

**Supplementary Figure 3: Bulk CB CD34+ HSPCs edited with three antibody cassettes simultaneously. a)** Allelic integration frequency at *CCR5* in CB CD34$^+$ HSPCs targeted with AAV6 cassettes for linker antibodies as indicated (n = 1 biological donor). **b)** Expected amplicon size for in-out PCR with primers designed to amplify each antibody individually when integrated in the *CCR5* locus. **c)** Gel electrophoresis analysis of amplicons from in-out PCR for integration of each antibody construct. Genomic DNA from CB CD34$^+$ HSPCs targeted as indicated for each lane was amplified simultaneously with all three sets of in-out PCR primers described in panel b.

**Supplementary Figure 3: Bulk CB CD34+ HSPCs edited with three antibody cassettes simultaneously**

**a**

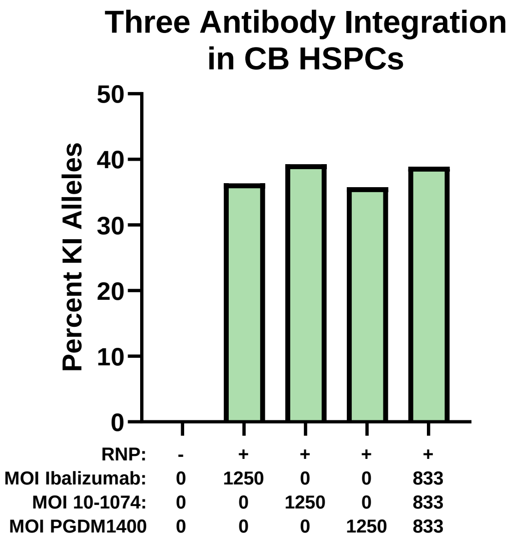


**b**

| In/Out PCR Primers | Expected Amplicon (bp) |
|---|---|
| Ibalizumab | 1498 |
| 10-1074 | 1761 |
| PGDM1400 | 2060 |

**c**

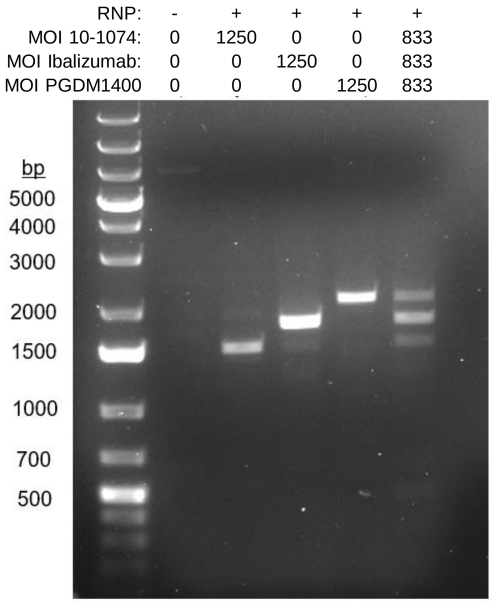

**Supplementary Figure 4: Maintenance of knockout and knock-in in xenografted NSG and NBSGW mice. a)** Distribution frequency of WT, INDEL, and KI human alleles in the bone marrow of NSG mice at 16 weeks post-transplantation (Fig. 3). Each number represents an individual mouse transplanted with cells targeted as indicated. **b)** Distribution frequency of WT, INDEL, and KI human alleles in the bone marrow of NBSGW mice at 12 weeks post-transplantation (Fig. 4). Each number represents an individual mouse transplanted with cells targeted as indicated. **c)** Percent of KI alleles from the bulk bone marrow integrated with 10-1074 or Ibalizumab within NBSGW mice receiving “10-1074+Ibalizumab” targeted cells (n = 5). Bars represent mean and errors bars represent SD.

**Supplementary Figure 4: Maintenance of knockout and knock-in in xenografted NSG and NBSGW mice**

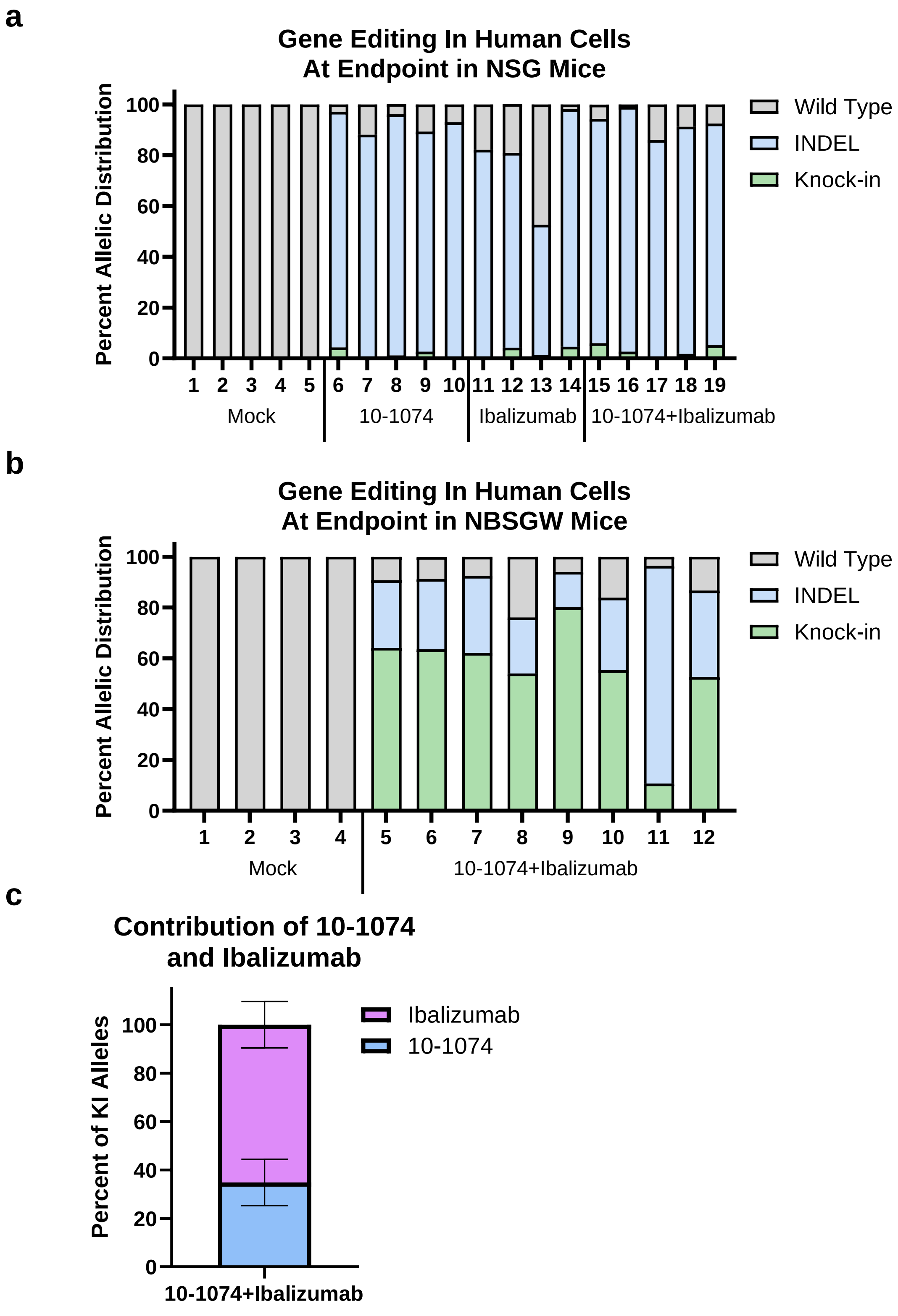

**Supplementary Figure 5: Human IgG in xenografted NSG and NBSGW mice. a)** Serum concentration of human IgG in NSG mice 16 weeks post-engraftment with CB CD34$^+$ HSPCs targeted as indicated and described in Fig. 3 (n = 4 for mock and Ibalizumab, n = 5 for 10-1074 and 10-1074+Ibalizumab). **b)** Serum concentration of human IgG in NBSGW mice 12 weeks post-engraftment with CB CD34$^+$ HSPCs targeted as indicated and described in Fig. 4 (n = 4 for mock, n = 8 for 10-1074+Ibalizumab). All dots represent individual mice engrafted with mock HSPCs or HSPCs edited with AAV6 and RNP for the linker antibody construct(s) indicated. All lines represent mean.

**Supplementary Figure 5: Human IgG in xenografted NSG and NBSGW mice**

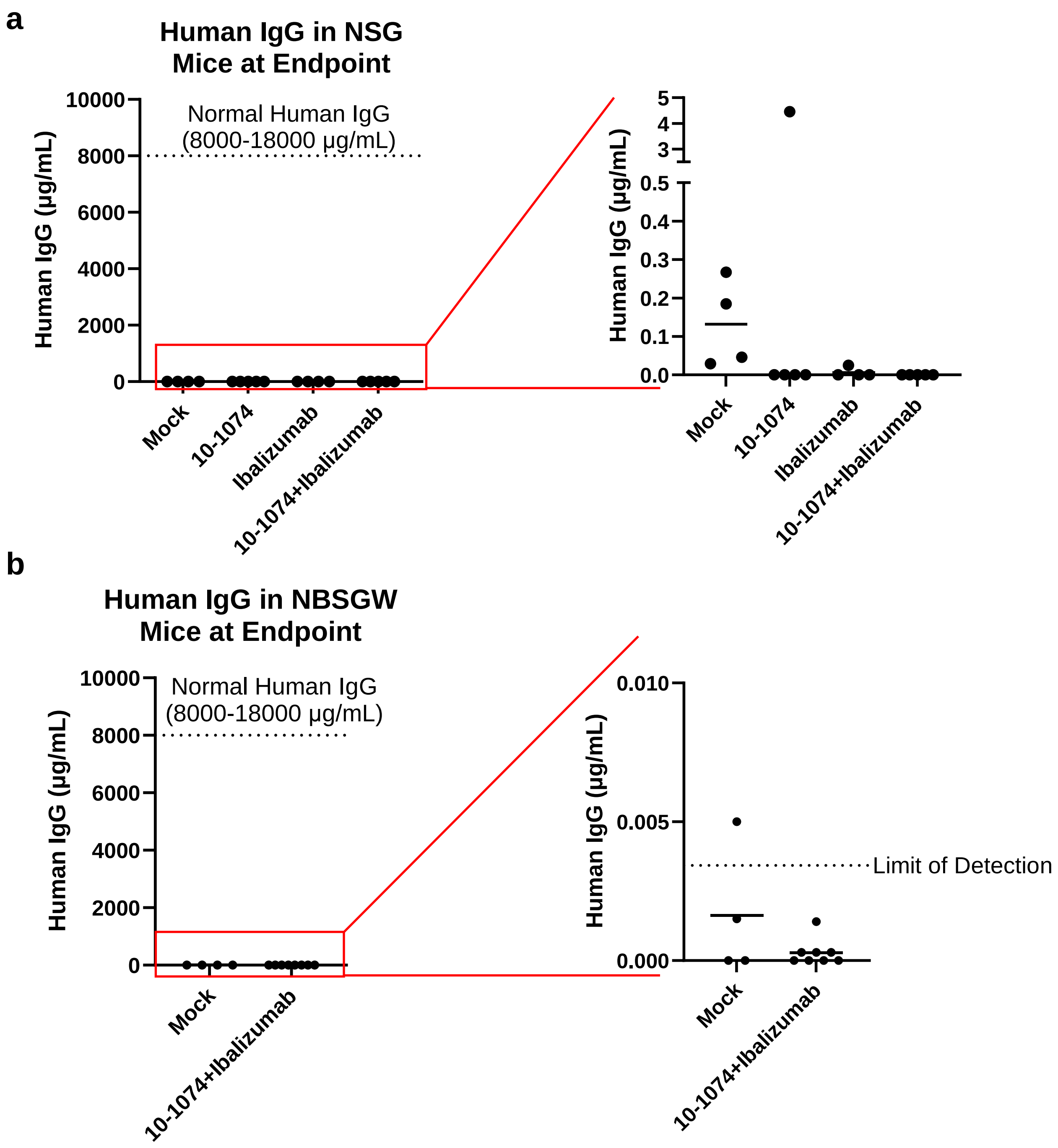

**Supplementary Figure 6: The EEK promoter is highly active in human B Cells. a)** Schematic for integration of cassettes with the EEK promoter (left) or the ubiquitous human UBC promoter (right) driving expression of GFP (EEK-GFP and UBC-GFP). UBC-GFP is integrated using a different *CCR5* sgRNA (*sg-CCR5-#2*) that cuts downstream within exon 2. **b)** Percent GFP+ cells from adult peripheral blood CD19+ B Cells targeted with AAV6 cassettes as indicated (n = 1 biological donor). Each AAV6 was used at an MOI of 25,000. **c)** Median fluorescence intensity of GFP+ B cells targeted as indicated (n = 1). Each AAV6 was used at an MOI of 25,000.

**Supplementary Figure 6: The EEK promoter is highly active in human B Cells**

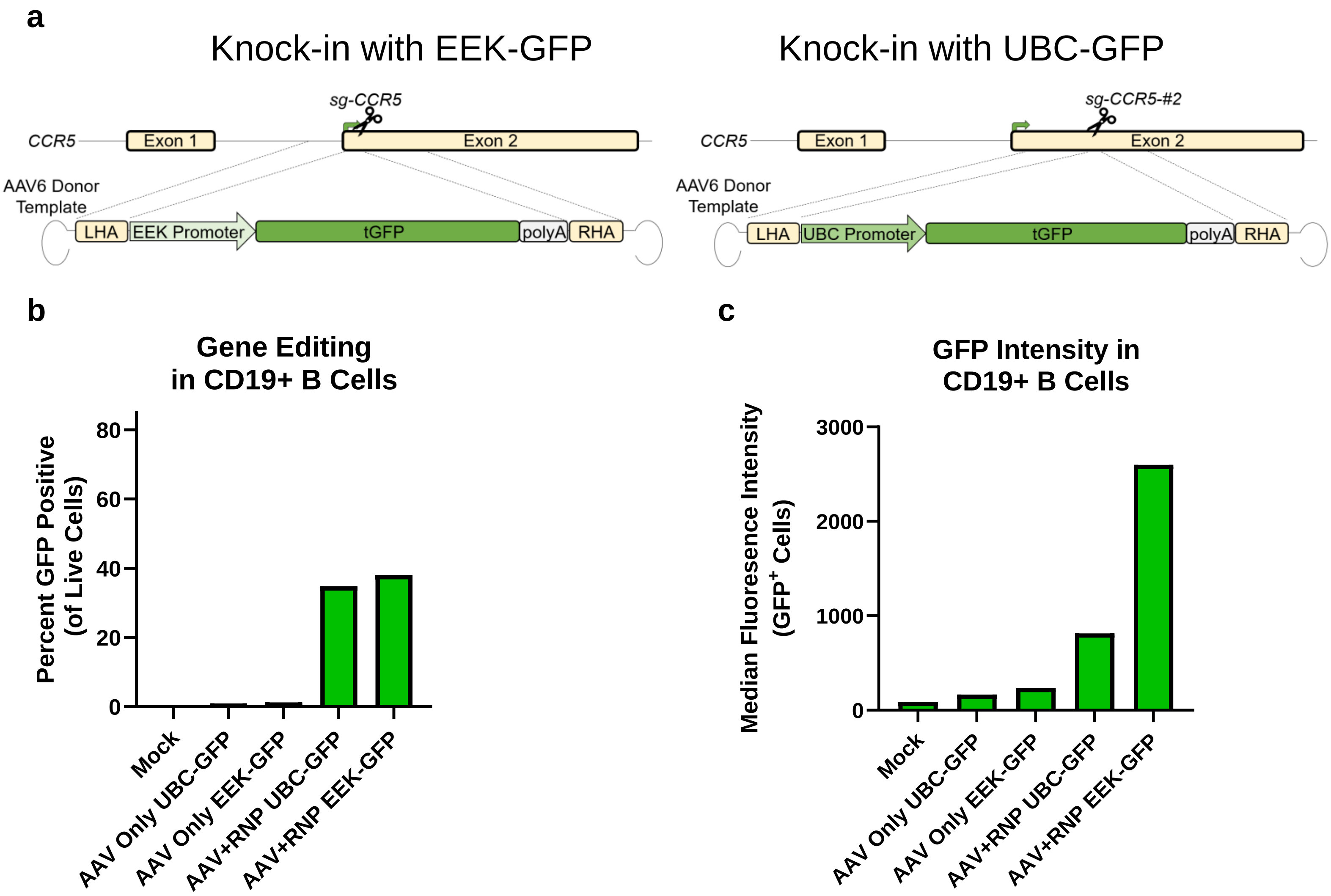

**Supplementary Figure 7: B cell subsets are not perturbed by expression of exogenous antibodies. a)** Representative gating strategy for classification of B cell subsets. Gating derived from Velounias and Tull, 2022.[91] **b)** Percent CD19+ positive cells from adult peripheral blood B Cells targeted with AAV6 cassettes as indicated (n = 1 biological donor). Each AAV6 was used at an MOI of 25,000. “EEK-10-1074” represents integration of linker 10-1074 corresponding Fig. 2A. Days are days post-editing. **c)** Distribution of cells within the CD19+ gate that are phenotypically plasmablasts (CD19+ CD27+ CD38+) or general B cells (CD19+ CD27+ CD38-, CD19+ CD27- CD38-, or CD19+ CD27- CD38+). **d)** Distribution of cells within the B cell gate that are phenotypically memory B cells (CD19+ CD27+ IgD-), marginal zone B cells (CD19+ CD27+ IgD+), Double negative B cells (CD19+ CD27- IgD-), or Naïve/Transitional B cells (CD19+ CD27- IgD-). **e)** Distribution of cells within the memory B cell gate that are phenotypically unswitched memory B cells (CD19+ CD27+ IgD- IgM+) or class switched memory B cells (CD19+ CD27+ IgD- IgM-).

**Supplementary Figure 7: B cell subsets are not perturbed by expression of exogenous antibodies**

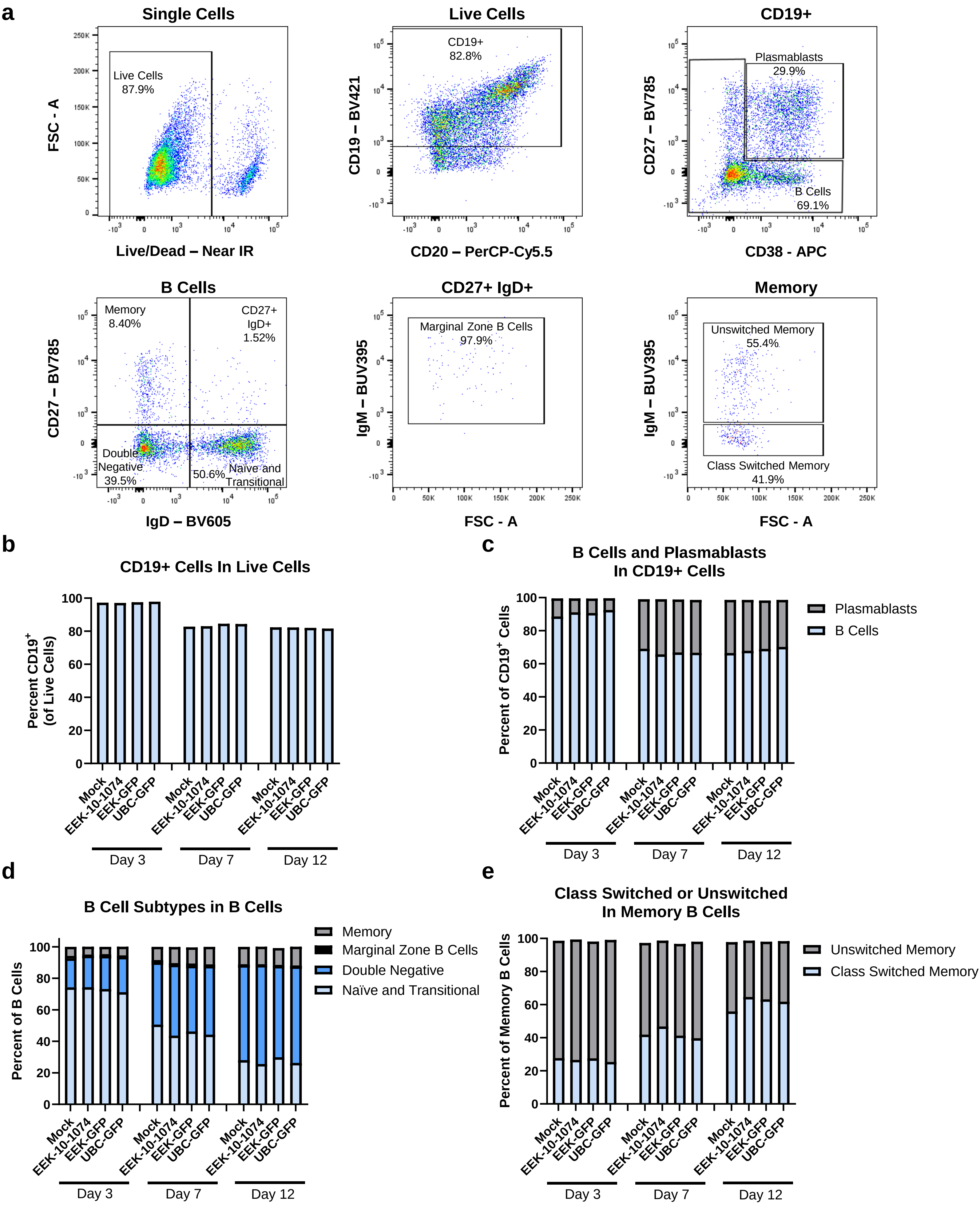

**Supplementary Figure 8: Engineered B cells secrete linker antibodies. a)** Concentration of each antibody as determined by antigen specific ELISA from B cell supernatant from cells targeted with AAV6 cassettes as indicated (n = 3 biological donors). Each AAV6 was used at an MOI of 25,000. B cells were plated at $1x10^6$ cells per mL and supernatant was collected after 5 days.

**Supplementary Figure 8: Engineered B cells secrete linker antibodies**

**a**

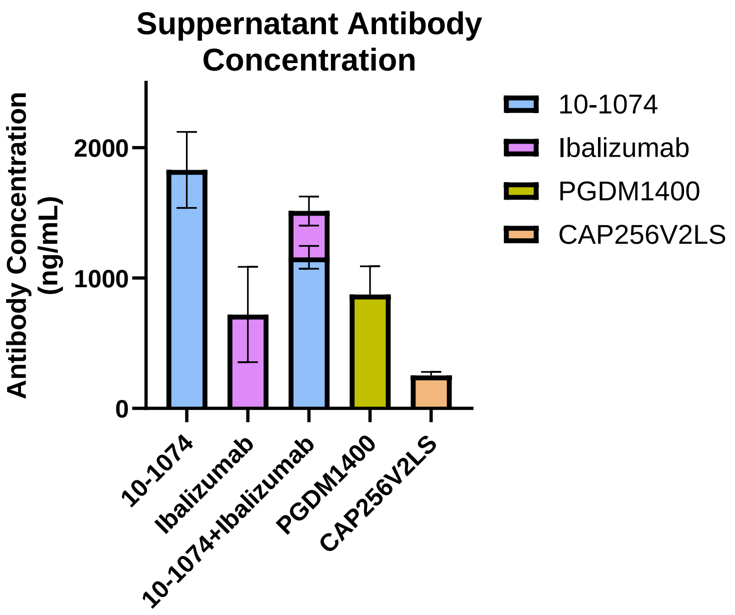
